# Plasma Ficolins Enable Liver Macrophages to Capture Blood-Borne Bacteria by Recognizing Capsular Polysaccharides

**DOI:** 10.64898/2025.12.22.696098

**Authors:** Xueting Huang, Haoze Chen, Jiaying Feng, Kunpeng Li, Jiao Hu, Yanhong Liu, Haoran An, Rui-Quan Zhou, Pei-Jun Yang, Hai-Dong Tan, Si-Yin Tan, Lok-To Sham, Ye Xiang, Jing-Ren Zhang

## Abstract

Ficolins are evolutionarily conserved glycan-binding lectins in plasma or on cell membranes of immune cells, yet their physiological functions remain to be defined. Here we demonstrate that plasma ficolins drives the capture of diverse encapsulated bacteria in the blood circulation by the liver resident macrophages (Kupffer cells). Ficolin-activated immunity in the liver sinusoids provides swift and sterilizing immunity in mice. Mice lacking ficolin-A (the only plasma ficolin) became hypersusceptible to the blood infections of the ficolin-A-recognizable bacterial serotypes. Mechanistically, plasma ficolins activate complement C3 on bacterial surfaces by pattern recognition of acetylated monosaccharides on polysaccharide capsules, and thereby deliver the opsonized bacteria to C3 receptors on Kupffer cells for phagocytic elimination. These findings reveal that plasma ficolins serve as pattern-recognition receptors that are essential for maintaining the bloodstream sterile.

## INTRODUCTION

Pattern recognition receptors (PRRs) are well-established for their ability to detect pathogen-associated molecular patterns (PAMPs), such as microbial glycans, flagella, lipids, and nucleic acids ^1, 2^. A variety of soluble and membrane-bound lectins play a critical role in recognizing microbial glycans, including capsular polysaccharides (CPS), lipopolysaccharides (LPS), cell wall peptidoglycans, and glycoproteins ^3^. Membrane-bound glycan receptors, such as Toll-like receptors (TLRs) and C-type lectin receptors (CLRs), are particularly well-documented for their importance in pathogen detection ^4–6^. In addition to membrane-bound receptors, a wide array of soluble lectins in the bloodstream, including ficolins, collectins, intelectins, mannose-binding lectins, and C-reactive proteins, also recognize microbial glycans ^7^. While molecular binding of these soluble lectins activates the complement system and thereby promotes opsonophagocytosis of pathogens under the *in vitro* conditions ^3, 7, 8^, it remains virtually unknown how soluble lectins contribute to pathogen clearance in blood infections, the deadliest form of microbial infections.

Since the discovery of the first ficolin in pig ^9^, ficolins are found in many vertebrates and invertebrates ^10^. Three ficolins are described in human: ficolin-1 (M-ficolin), ficolin-2 (L-ficolin), and ficolin-3 (H-ficolin). Mice possess ficolin-A (FCN-A) and ficolin-B (FCN-B), with the third ficolin gene as a pseudogene ^11^. FCN-A and the human ortholog ficolin-2 are mainly synthesized in the liver and present in plasma; FCN-B is more closely related to human ficolin-1 with a dominant expression by myeloid cells ^12, 13^. All ficolins consist of the *N*-terminal collagen-like domain and C-terminal fibrinogen-like domain ^14^. While the former is required for ficolin oligomerization, the fibrinogen-like domain recognizes the acetyl group of microbial carbohydrates ^15–17^. In particular, human ficolin-2 recognizes capsular polysaccharides of *Streptococcus pneumoniae* ^18–22^, *Streptococcus agalactiae* ^23–25^ and *Staphylococcus aureus* ^18^. Mouse FCN-A binds to multiple bacteria and bacterial cell wall components *in vitro* ^17, 26^, whereas FCN-B recognizes sialic acid ^17^. Mice lacking ficolin-A and -B show a modest increase in susceptibility to serotype-2 *S. pneumonia*e infection ^27^. Many *in vitro* studies have demonstrated that the ficolin binding to bacterial glycans activates the complement lectin pathway and thereby promotes opsonophagocytosis ^17, 28–30^. However, it remains unclear whether ficolins contribute to host defense against microbial infections under physiological conditions.

Invasive infections by encapsulated bacteria are the leading causes of septic infections, the deadliest form of infections. Capsules are the outmost structures of many bacteria. A recent study has identified the capsule biosynthesis genes in more than 50% of bacterial genomes, indicating the high prevalence of the capsules in the prokaryotes ^31^. The capsules are important virulence factors in many pathogenic bacteria, which protect bacteria from being detected by immune molecules and cells ^32, 33^. Our recent studies have revealed that the capsules promote bacterial survival and thereby virulence mainly by fending off the pathogen recognition and capture by liver-resident macrophages - Kupffer cells (KCs) during blood infections ^34, 35^. However, the immune evasion potency of the capsules greatly varies depending on capsular structures or serotypes. While a small fraction of capsule types fully protects bacteria from host recognition during blood infections, the vast majority of capsules are still recognizable by the capsule-binding receptors. However, only a few capsule receptors have been identified to promote the hepatic clearance of encapsulated bacteria, including the asialoglycoprotein receptor (ASGR) ^34^, natural antibodies ^36^, and C-reactive protein (CRP) ^37^.

In this study, we demonstrate that plasma ficolins from humans (ficolin-2) and mice (ficolin-A) act as PRRs, enabling KCs to capture a broad range of encapsulated bacteria. Using ficolin-A-deficient mice, we show that ficolins play essential roles in maintaining blood sterility and host survival. Comparative analysis reveals notable differences in ligand recognition between human and mouse ficolin orthologs. We finally discuss the implications of ficolin-mediated bacterial clearance for innate immunity and host specificity.

## RESULTS

### Ficolin-A binds to serotype-19F capsule of *S. pneumoniae*

Serotype-19F *S. pneumoniae* (*Spn-*19F) is one of the most prevalent serotypes in childhood invasive pneumococcal disease (IPD) cases in many countries before the introduction of pneumococcal conjugate vaccines ^38, 39^. However, mice are highly resistant to *Spn-*19F infection ^34^. *Spn-*19F bacteria are rapidly cleared from the bloodstream by Kupffer cells (KCs) in the liver sinusoids of mice, which is serotype-specifically blocked by *Spn-*19F capsular polysaccharide (CPS19F) ^34^. This finding indicated that mouse has a specific receptor(s) for CPS19F to mediate hepatic capture of *Spn-*19F. CPS19F-coupled beads were incubated with membrane proteins from mouse liver nonparenchymal cells in the presence of 20% mouse serum (**Fig. 1A**). Mass spectrometry analysis revealed a total of 676 proteins significantly enriched by CPS19F, as compared with serotype-8 CPS (CPS8) (≥2 folds) (**Fig. 1B**; **Table S1**). CPS8-conjugated beads were used as a negative control, due to the poor hepatic capture of serotype-8 pneumococci ^34^.

**Figure 1.**
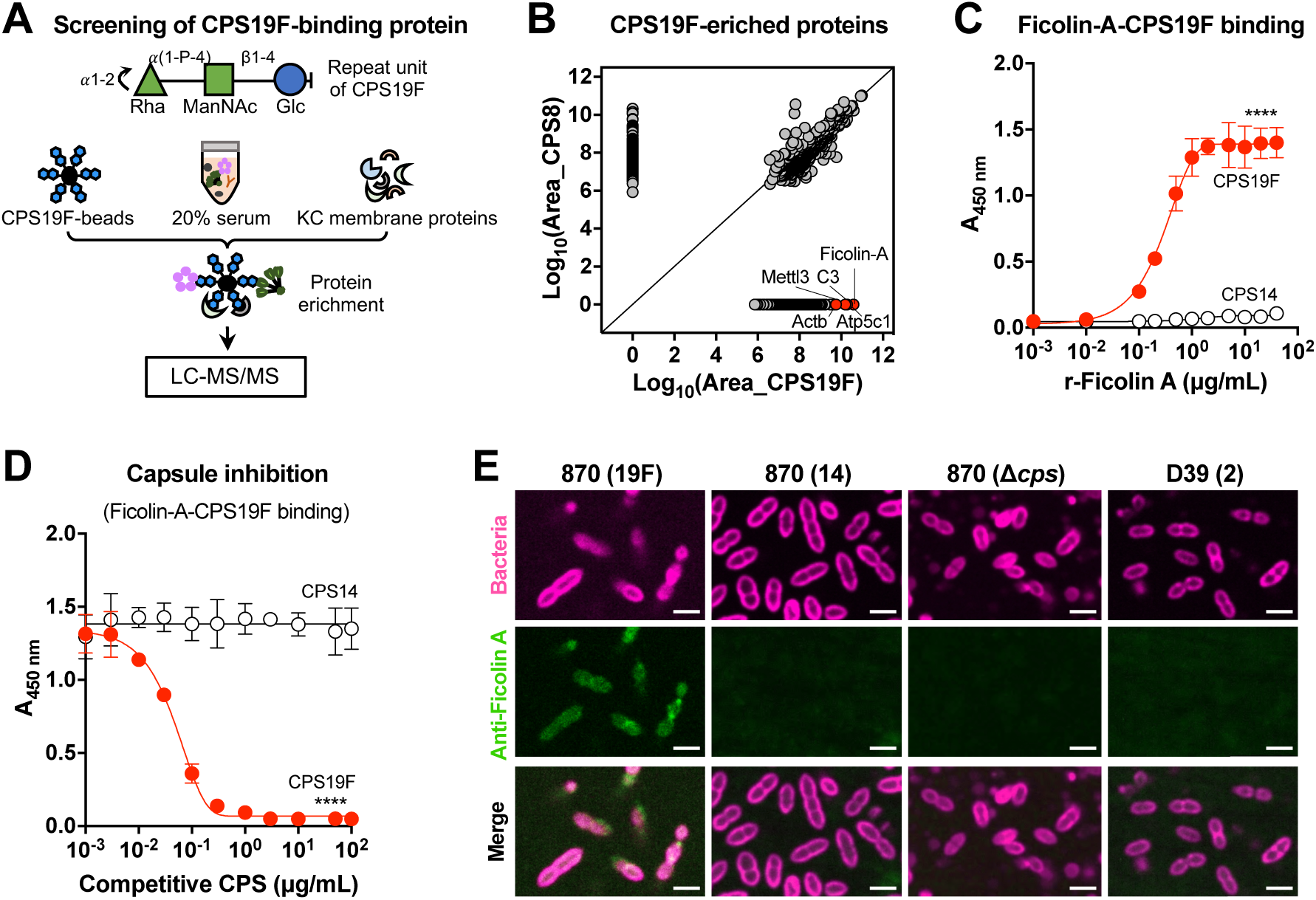
Specific binding of ficolin-A to serotype-19F capsule. **A.** Schematic illustration of experimental approach to identify CPS19F-binding proteins. The repeat unit of CPS19F are shown at the top. **B.** CPS19F-enriched proteins are plotted against the proteins pulled down by CPS8-conjugated beads. The top 5 hits are indicated as red circles. **C.** Ficolin-A (FCN-A) binding to CPS19F was assessed by ELISA using CPS19F- or CPS14 (negative control)-coated wells and recombinant FCN-A (r-FCN-A); the level of CPS19F binding presented as the absorbance at 450 nm. n = 3. **D.** Competitive inhibition of rFCN-A-CPS19F binding interaction by free CPS19F. rFCN-A was incubated with free CPS19F before being added to CPS19F-coated wells for ELISA as in (C). n = 3. **E.** Specific binding of ficolin-A to serotype-19F pneumococci. Isogenic ST870 derivatives each producing a unique capsule or no capsule (Δ*cps*) were pre-labelled by AF647 (pink) and incubated with mouse serum, and visualized by immunofluorescence microscopy after being sequentially stained with anti-FCN-A and FITC-conjugated secondary antibodies (green). Scale bar, 2 μm. These data represent the results of at least two independently experiments. The data in C and D are presented as mean ± SD, with statistical differences determined by Two-way ANOVA test, \*\*\*\**P* < 0.0001.

Ficolin-A (FCN-A) was found to be the most abundantly enriched protein (**Fig. 1B**), one of the two known ficolin proteins in mouse. FCN-A is a liver-derived plasma protein of 334 amino acids with a concentration of approximately 3.5 µg/mL in serum ^13, 40^. We further characterized FCN-A interaction with CPS19F using a recombinant FCN-A (rFCN-A). The results revealed a dose-dependent binding of rFCN-A with purified CPS19F. In contrast, rFCN-A did not show obvious binding to CPS of serotype-14 pneumococci (CPS14), which is recognized by the asialoglycoprotein receptor (ASGR) ^34^ (**Fig. 1C**). In a consistent pattern, free CPS19F but not CPS14 competitively blocked the rFCN-A binding to immobilized CPS19F in a dose-dependent manner (**Fig. 1D**), confirming that ficolin-A specifically recognizes CPS19F.

We further tested specific binding of ficolin-A to the intact capsule of *Spn-*19F. Immunofluorescence imaging revealed that rFCN-A bound to *Spn-*19F (TH870^19F^), but not isogenic bacteria producing a serotype-14 capsule (TH870^14^) (**Fig. 1E**). We next assessed potential interaction of FCN-A with the cell wall of *S. pneumoniae,* based on the previous report that ficolin-2, the major plasma form of human ficolins, recognizes lipoteichoic acid, a cell wall component of Gram-positive bacteria ^41, 42^. The acapsular pneumococci (TH870Δ*cps*) did not show obvious binding to rFCN-A (**Fig. 1E**). Endo *et al.* described that the FCN-A-deficient mice are more susceptible to the infection of serotype-2 *S. pneumoniae* (strain D39). However, consistent with the full evasion of hepatic capture by the serotype-2 capsule ^34^, rFCN-A did not show obvious binding to the intact D39 bacteria.

### Ficolin-A enables liver macrophages to capture blood-borne serotype-19F pneumococci

To assess the role of FCN-A in host defense, we tested the impact of genetic deficiency on the hepatic capture of blood-borne *Spn-*19F. *Spn-*19F bacteria were rapidly cleared to an undetectable level in the bloodstream of wildtype (WT) mice (<500 CFU/mL) in the first 20 min post intravenous (i.v.) inoculation. In contrast, FCN-A-deficient (*Fcna*^-/-^) mice completely failed to clear the bacteria (**Fig. 2A**). The 50% clearance time (CT_50_) of *Spn-*19F was extended from 1.2 min in WT mice to >30 min in *Fcna*^-/-^ mice (**Table S2**). However, *Spn-*14 bacteria were similarly cleared from the bloodstream of WT (CT_50_ = 0.4 min) and *Fcna*^-/-^ mice (CT_50_ = 0.5 min). This result showed that ficolin-A is essential for capsule type-specific clearance of *Spn-*19F from the blood circulation in mice.

**Figure 2.**
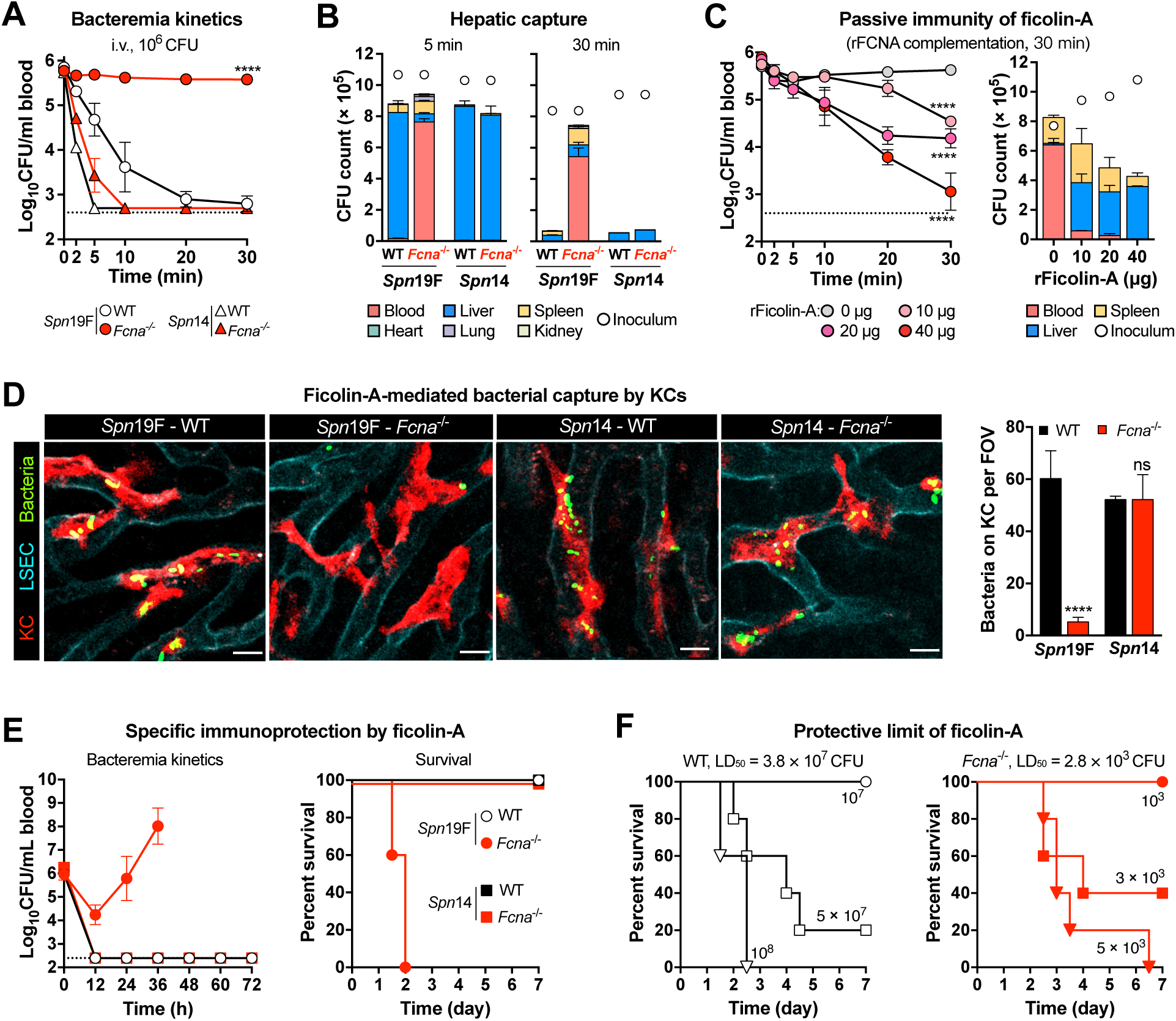
Ficolin-A-activated capture of *Spn-*19F by Kupffer cells (KCs) in the liver sinusoids. **A.** The role of FCN-A in the *Spn-*19F clearance was assessed by blood CFU plating of WT and ficolin-A-deficient (*Fcna*^-/-^) mice post i.v. infection with 10^6^ CFU of *Spn-*19F or *Spn-*14 (negative control). n = 3. Dotted line indicates the detection limit. **B.** The impact of FCN-A deficiency on hepatic clearance of *Spn-*19F in *Fcna*^-/-^ mice, in terms of bacterial level in blood and five major organs at 5 and 30 min post infection as in (A). The inoculum is indicated with an open circle at the top. n = 3. **C.** Complementation of the immuno-deficiency of *Fcna*^-/-^ mice by i.v administration of rFCN-A 2 min before infection as in (A). n = 3. **D.** Capture of bacteria (green) by KCs (red) was visualized by IVM in the context of liver vasculatures (cyan) of WT and *Fcna*^-/-^ mice post i.v. infection with 5×10^7^ CFU of *Spn-*19F or *Spn-*14 (negative control). KC-associated bacteria are presented as immobilized bacteria per field of view (FOV). Scale bar, 10 μm. n = 2. **E.** FCN-A-mediated protection against *Spn-*19F infection is presented as bacteremia (left) and the survival rate of infected mice (right) post i.v. infection with 10^6^ CFU. n = 5. **F.** Protective potency of FCN-A against *Spn-*19F was characterized by comparing survival rate of WT (left) and *Fcna*^-/-^ mice post i.v. infected with various doses of *Spn-*19F. The LD_50_ values are shown at the top. n = 5. The representative data are presented as mean ± SD, with statistical differences determined by Two-way ANOVA (A and C), *t* tests (D, right panel) and log-rank test (E and F). *, *P* < 0.05; **, *P* < 0.01; ****, *P* < 0.0001; ns, no significant difference.

Our previous study showed that the liver is the major immune organ to trap and kill blood-borne *Spn-*19F ^34^. We thus tested the impact of FCN-A deficiency on hepatic capture of *Spn-*19F. While the vast majority of the *Spn-*19F inoculum was found in the liver of WT mice at 5 min post i.v. infection (mpi), only there was a marginal level of hepatic bacteria in *Fcna*^-/-^ mice at this time point (**Fig. 2B**). This result explains the great difference in blood bacteria between WT and *Fcna*^-/-^ mice at 5 min (**Fig. 2A**). At 30 mpi, WT mice nearly completely eliminated *Spn-*19F pneumococci from the liver, but the bacteria were still retained in the bloodstream of *Fcna*^-/-^ mice (**Fig. 2B**, right panel). In contrast, *Spn-*14 bacteria were equally captured in the livers of *Fcna*^-/-^ and WT mice at 5 mpi and eradicated at 30 mpi. The FCN-A-mediated immunity against *Spn-*19F was further verified by passive i.v. administration of rFCN-A prior to infection. The treatment with rFCN-A enabled *Fcna*^-/-^ mice to shuffle *Spn-*19F from the bloodstream to the liver in a dose-dependent fashion (**Fig. 2C**). Collectively, these data showed that ficolin-A drives the hepatic capture of *Spn-*19F.

To characterize the cellular basis of ficolin-triggered bacterial capture in the liver, we visualized bacterial traffic in the sinusoidal vasculatures by intravital microscopy (IVM) imaging. Once entering the blood vessels in the livers of WT mice, *Spn-*19F bacteria were rapidly tethered to the endothelium-embedded Kupffer cells (KCs) of WT mice (**Fig. 2D**; **Videos S1** and **S2**). However, *Spn-*19F pneumococci freely passed by KCs in the liver sinusoids of *Fcna*^-/-^ mice. As a negative control, *Spn-*14 bacteria were similarly captured by KCs of *Fcna*^-/-^ and WT mice. This information verified KCs are the primary immune cells that are responsible for FCN-A-mediated bacterial capture in the liver. The FCN-A-mediated KC capture of *Spn-*19F bacteria was also validated *in vitro*. Primary mouse KCs showed substantial capture of *Spn-*19F bacteria in the presence of normal serum, but not FCN-A-deficient serum (**Fig. S1A**). Exogenous supplementation of FCN-A-deficient serum with rFCN-A significantly enhanced *Spn-*19F adhesion. However, the presence of serum did not significantly affect KC capture of *Spn-*14 pneumococci. These data demonstrated that FCN-A is essential for specific capture of serotype-19 pneumococci by KCs in the liver.

To assess the immuno-protectivity of FCN-A-mediated bacterial capture in the liver, we compared the survival rate between WT and *Fcna*^-/-^ mice post *Spn-*19F infection. WT mice completely cleared *Spn-*19F bacteria from the bloodstream in the first 12 hr post infection, and survived the infection (**Fig. 2E**). *Fcna*^-/-^ mice significantly cleared *Spn-*19F, which was likely accomplished by the splenic red pulp macrophages ^43^. However, the KO mice displayed worsening bacteremia at the later stage of infection and succumbed to the infection within 48 hr post infection. By comparison, *Fcna*^-/-^ mice did not show obvious deficiency against serotype-14 pneumococci, in terms of bacteremia and survival. Further experiments uncovered a 13,571-fold difference in the median lethal dose (LD_50_) of *Spn-*19F infection between WT (3. 8×10^7^ CFU) and *Fcna*^-/-^ (2.8×10^3^ CFU) mice (**Figs. 2F** and **S1B**). In contrast, mice lacking ficolin-B (*Fcnb*^-/-^), the other mouse ficolin (**Fig. S1C**), did not show significant impairment against *Spn-*19F (**Fig. S1D**), which is consistent with the fact that FCN-B is mainly expressed as a cellular form in the bone marrow and barely detectable in the blood ^44^. These data demonstrated that the capsule recognition by FCN-A represents a robust anti-bacterial immunity.

### Ficolin-A broadly recognizes multiple pneumococcal capsule types

To understand if ficolin-A also recognizes other capsular variants of *S. pneumoniae* beyond serotype-19F, we tested the impact of FCN-A deficiency on the early clearance of 69 additional serotypes in our collection, including 10 of the 14 serotypes that are shown to bind to human ficolin-2 (FCN-2), the mouse ficolin-A ortholog (11A, 11D, 15F, 31, 33A, 35A, 35C, 42, 47A and 47F) ^18–20^. This screen identified 8 additional serotypes whose clearance from the bloodstream was significantly impaired in *Fcna*^-/-^ mice (9A, 9L, 9N, 9V, 19A, 19B, 19C and 18A) (**Figs. 3A** and **S2A**; **Table S2**). These serotypes are collectively referred to as “ficolin-sensitive” serotypes. While the FCN-A deficiency made the host fully lose the immunity in clearing all the ficolin-sensitive serotypes, except for serotype 19B, indicating ficolin-A is a dominant receptor recognizing CPSs of these pneumococci. The partial clearance of serotype-19B bacteria was likely due to the operation of the other receptor(s). Except for 18A, all of the ficolin-sensitive serotypes belong to serogroups 9 and 19. Interestingly, all of the 10 FCN-2-binding serotypes were normally cleared from the circulation of *Fcna*^-/-^ mice (**Table S2**), indicative of distinct ligand specificities between the human and mouse ficolins.

**Figure 3.**
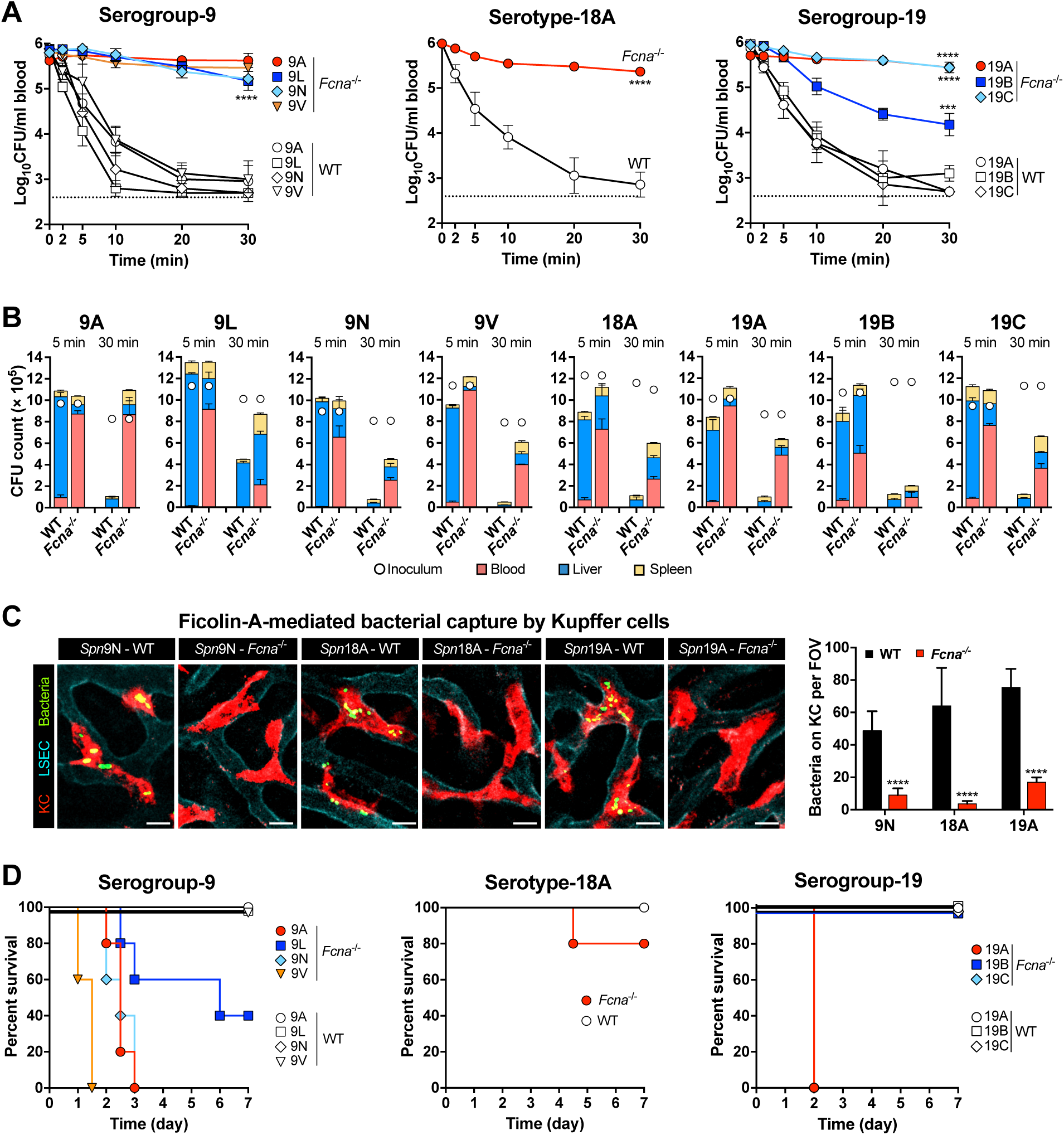
Ficolin-A-mediated broad immunity against diverse pneumococcal serotypes. **A.** Bacteremia kinetics of *Fcna*^-/-^ and WT mice post i.v. infection with 10^6^ CFU of serogroup-9, serotype-18A or serogroup-19 pneumococci. n = 3. **B.** Bacterial capture (5 min) and killing (30 min) in the liver post i.v. infection with 10^6^ CFU of serogroup-9, serotype-18A or serogroup-19 pneumococci. n = 3. **C.** IVM imaging of KC capture of serogroup-9, serotype-18A and serogroup-19 pneumococci in *Fcna*^-/-^ and WT mice post i.v. infection 5 × 10^7^ CFU. n = 2. **D.** Survival of *Fcna*^-/-^ and WT mice post i.v. infection with 10^6^ CFU of serogroup-9, serotype-18A or serogroup-19 pneumococci. n=5. The representative data are presented as mean ± SD, with statistical differences determined by Two-way ANOVA (A), *t* tests (C, right panel) and log-rank test (D). *, *P* < 0.05; **, *P* < 0.01; ***, *P* < 0.001; ****, *P* < 0.0001; ns, no significant difference.

The importance of FCN-A in serotype-specific immunity was confirmed by the loss of hepatic capture/killing of FCN-A-sensitive serotypes in *Fcna*^-/-^ mice (**Fig. 3B**). While the vast majority of the eight pneumococcal serotypes were shuffled from the blood circulation to the livers of WT mice at 5 mpi, the same inoculum mostly persisted in the blood of *Fcna*^-/-^ mice with marginal levels of hepatic bacteria. At 30 mpi, WT mice infected with the FCN-A-sensitive serotypes showed dramatic reduction in total bacterial burden in the livers, demonstrating rapid killing of liver-trapped bacteria. In sharp contrast, *Fcna*^-/-^ mice still carried much more bacteria for all the eight FCN-A-sensitive serotypes, which were mostly in the blood circulation. Similar to their variations in the early clearance from the blood circulation of *Fcna*^-/-^ mice, FCN-A-sensitive serotypes showed substantial differences in hepatic capture/killing. In particular, serotype-9A bacteria were completely maintained at the inoculation levels at 30 mpi in *Fcna*^-/-^ mice, indicating that FCN-A is the only capsule-binding receptor for these serotypes. In contrast, the levels of the other seven serotypes (9L, 9N, 9V, 18A, 19A, 19B and 19C) were still significantly reduced in *Fcna*^-/-^ mice at 30 mpi. This variation supports the operation of the other redundant receptor(s) to these capsules.

Consistent with the dominant role of liver-resident macrophages in capturing serotype-19F pneumococci (**Fig. 2D**), IVM imaging revealed that KCs effectively captured FCN-A-sensitive serotypes 9N, 18A and 19A in the liver sinusoids of WT mice, but these serotypes freely passed by KCs of *Fcna*^-/-^ mice in the liver sinusoids (**Fig. 3C**; **Videos 3-5**). Consistent with relatively milder impact of FCN-A deficiency on the early clearance and hepatic capture of serotype-19B pneumococci (**Figs. 3A** and **3B**), KCs of *Fcna*^-/-^ mice showed relatively less impairment in capturing these bacteria. This result demonstrated that ficolin-A broadly promotes the capsule type-specific clearance of blood-borne pneumococci by enabling KCs to capture these bacteria in the liver sinusoids.

Blood infection experiments further demonstrated the broad immunity of FCN-A against invasive infection of FCN-A-sensitive pneumococcal serotypes. *Fcna*^-/-^ mice were hyper-susceptible to infection of serotype-9A, -9N, -9V and -19A pneumococci (**Figs. 3D** and **S2B-D**). FCN-A deficiency yielded less severe impact on host defense against serotype-9L and -18A pneumococci, with partial mortality post i.v infection with 10^6^ CFU. All of *Fcna*^-/-^ mice survived septic infection with 10^6^ CFU of serotype-19B or -19C pneumococci, indicating the operation of FCN-A-independent hepatic clearance mechanisms against these serotypes. Collectively, these results demonstrated that ficolin-A is a broad-spectrum capsule receptor for at least 9 pneumococcal serotypes.

### Ficolin-A recognizes the capsules of multiple Gram-negative pathogens

We next determined potential contribution of ficolin-A to the clearance of encapsulated Gram-negative bacteria by assessing the impact of genetic deficiency on the clearance of *Haemophilus influenzae* and *Escherichia coli* strains in our collection (**Table S2**). As compared with bacteremia dynamics of WT mice in the first 30 mpi, *Fcna*^-/-^ mice completely lost the ability to clear two of the six *H. influenzae* serotypes. WT mice effectively cleared all the *H. influenzae* serotypes except for serotype-e (*Hi-*e) (**Fig. S3A**). By comparison, *Fcna*^-/-^ mice lost this immunity against serotype-c (*Hi-*c) (**Fig. 4A**, left panel) and serotype-f (*Hi-*f) (**Fig. 4A**, middle panel). The majority of *Hi-*c and *Hi-*f bacteria remained in the bloodstream of the KO mice in the first 30 mpi. This result was verified by significant impairment of *Fcna*^-/-^ mice in hepatic capture of *Hi-*c and *Hi-*f. Out of four *E. coli* serotypes with the rapid clearance in WT mice (**Fig. S3A**), the clearance of serotype KG2-1 was significantly retarded in the absence of ficolin-A (**Fig. 4A**, right panel). These results demonstrated that ficolin-A mediates the serotype-specific clearance of Gram-negative bacteria in the livers of mice.

**Figure 4.**
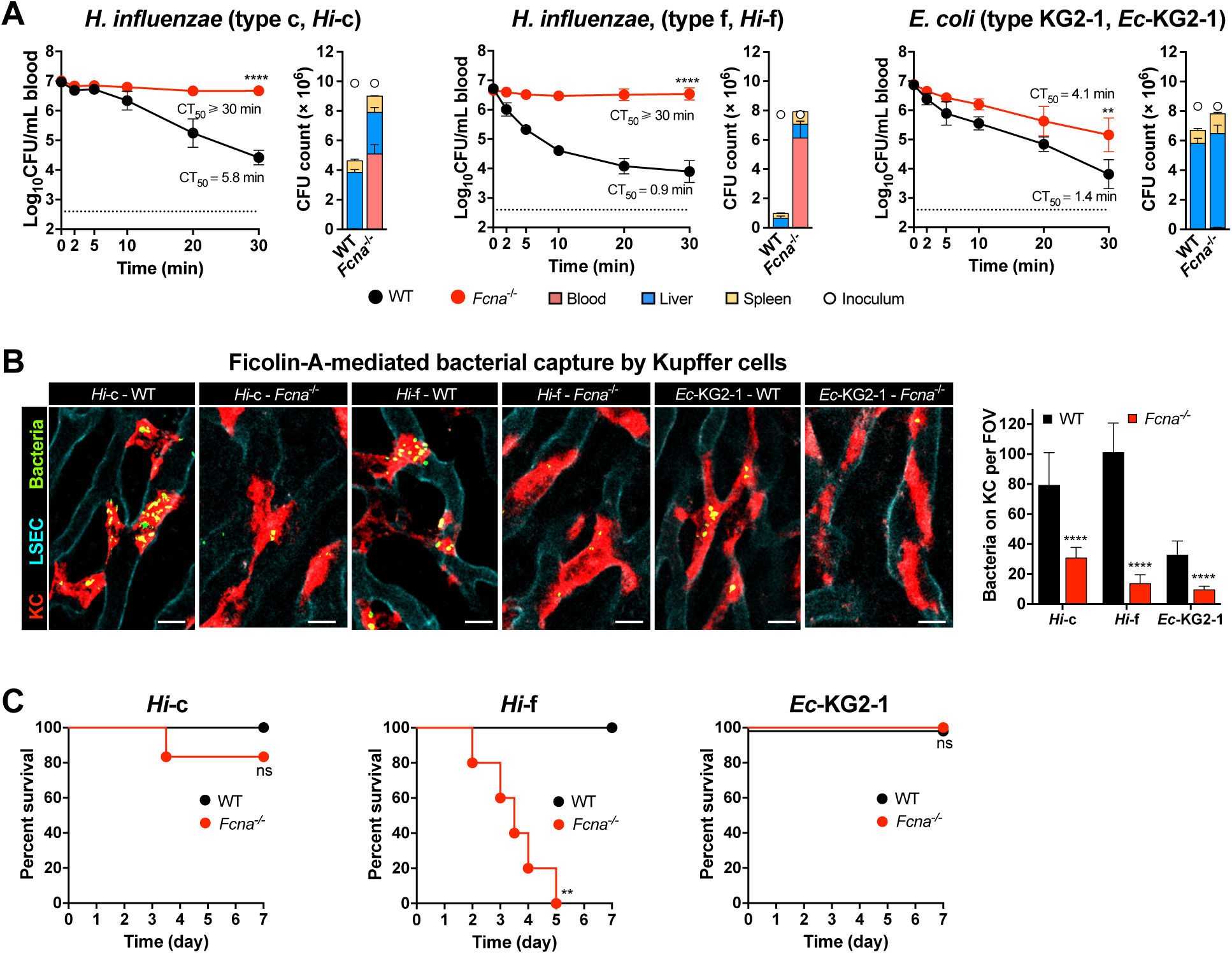
Ficolin-A-mediated serotype-dependent clearance of Gram-negative pathogens. **A.** The contribution of FCN-A to the clearance from bloodstream and hepatic capture of serotype*-*c (left) and serotype*-*f (middle) of *H. influenzae*, and serotype KG2-1 of *E. coli* (right) was characterized by CFU plating of the blood and major organs of *Fcna*^-/-^ and WT mice after i.v. infection with 10^7^ CFU. The CT_50_ value is presented for each serotype. n = 3-4. **B.** FCN-A-mediated capture of *H. influenzae* and *E. coli* by KCs in the liver sinusoids of *Fcna*^-/-^ and WT mice was imaged and quantified as in 2D. n = 2. **C.** FCN-A-based immunity against *Hi-*c (left), *Hi-*f (middle) and *E. coli* KG2-1(right) was assessed post i.v. infection with 10^8^ CFU as in 2E. n = 5. The representative data are presented as mean ± SD, and the statistical differences were determined by Two-way ANOVA with Tukey’s multiple comparisons test (A), multiple *t* tests (B, right panel) and log-rank test (C). **, *P* < 0.01; ****, *P* < 0.0001; ns, no significant difference. Shishi.

As observed with *Spn-*19F and other FCN-A-sensitive *S. pneumoniae* serotypes, IVM imaging showed rapid capture of *Hi-*c and *Hi-*f by KCs, but KCs-immobilized bacteria were significantly reduced in *Fcna*^-/-^ mice (**Fig. 4B**; **Videos 6, 7**). Consistent with greater contribution of FCN-A to the early clearance of *Hi-*f than *Hi-*c (**Figs. 4A**, left and middle panel), the liver macrophages of *Fcna*^-/-^mice showed more severe impairment in capturing serotype-*f H. influenzae*. In a similar manner, the level of KC-immobilized *E. coli* KG2-1 was significantly reduced in *Fcna*^-/-^ mice (**Fig. 4B**; **Video 8**). These imaging data demonstrated that FCN-A promotes serotype-specific hepatic capture of *H. influenzae* and *E. coli*.

We finally assessed the contribution of FCN-A to host resistance to blood infection of Gram-negative bacteria. Consistent with relatively more severe impact of FCN-A deficiency on the hepatic capture of *Hi-*f, all *Fcna*^-/-^ mice succumbed to i.v. infection of *Hi-*f as compared with partial mortality of the *Hi-*c*-*infected group (**Figs. 4C** and **S3B**). FCN-A deficiency did not show significant impact on host immunity against *E. coli* KG2-1 under the infection conditions, which agrees with the modest impact of ficolin-A deficiency on the clearance of *E. coli* KG2-1 (**Fig. 4A**). In summary, these results have revealed that ficolin-A confers capsule type-specific immunity to invasive infections of encapsulated Gram-negative bacteria.

### Ficolin-A-triggered bacterial capture in the liver requires the complement lectin pathway

Human ficolin-2 activates the complement system to promote bacterial opsonophagocytosis upon binding to microbial ligands ^45, 46^. We thus assessed the early clearance of *Spn-*19F in C3-deficient mice, based on the essential role of C3 in the complement-mediated opsonophagocytosis ^47^. *Spn-*19F clearance was significantly retarded in *C3*^-/-^ mice, in terms of bacteremia kinetics and bacterial capture in the liver (**Figs. 5A**, left panel; **S4A**). Under the *in vitro* conditions, FCN-A deficiency impaired the C3 deposition on CPS19F (**Fig. S4B**). These data suggest that FCN-A binding to *Spn-*19F promotes bacterial clearance by activating the C3 deposition on capsules. It should be noted that *C3*^-/-^ mice showed a partial deficiency as compared with the complete loss of *Spn-*19F clearance in *Fcna*^-/-^ mice (**Fig. 2A**). This finding suggests that FCN-A also engages a complement-independent mechanism(s) to shuffle blood-borne bacteria to KCs. However, *C3*^-/-^ mice phenotypically behaved as *Fcna*^-/-^ mice in bacterial clearance when the infection dose was increased to 10^7^ CFU (**Fig. 5A**, right panel). Pandra *et al.* reported that natural antibodies promote Fc receptor-dependent opsonophagocytosis of *Pseudomonas aeruginosa* by binding to ficolin-opsonized bacteria^48^. However, mice lacking both C3 and antibodies (μMT/*C3*^-/-^) showed the same pattern of bacterial clearance as *C3*^-/-^ mice (**Fig. S4C**), excluding the role of natural antibodies in FCN-A-initiated bacterial capture by KCs. Together, these results revealed that ficolin-A engages both the C3-dependent and -independent mechanisms for bacterial clearance.

**Figure 5.**
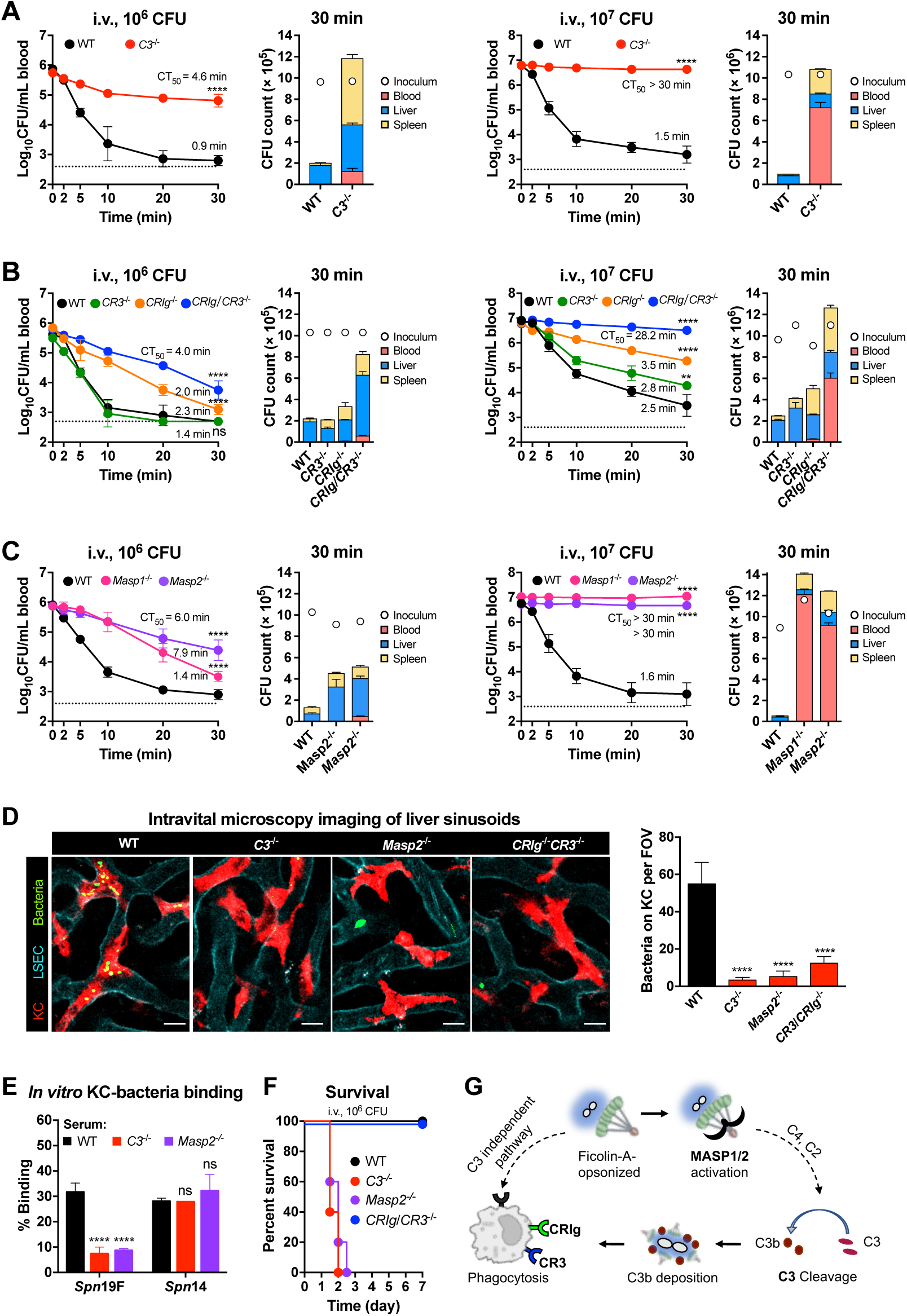
The requirement of the complement system for Ficolin-A-based immunity. **A.** The contribution of C3 to hepatic clearance of *Spn-19F*. Bacteria in the blood (left) and liver/spleen (right) of *C3*^-/-^ mice were quantified post i.v. infection with 10^6^ (left) or 10^7^ CFU (right). n = 3. **B.** The role of C3 receptors to hepatic clearance of *Spn-19F*. *CRIg*^-/-^, *CR3*^-/-^ and *CRIg*/*CR3*^-/-^ mice were infected with 10^6^ (left) or 10^7^ CFU (right) to quantify bacteria as in (A). n = 3. **C.** The role of the lectin pathway in hepatic clearance of *Spn-19F*. *Masp1*^-/-^ and *Masp2*^-/-^ mice were infected with 10^6^ (left) or 10^7^ CFU (right) to quantify bacteria as in (A). n = 3. **D.** Visualization of bacterial capture by KCs of complement-deficient mice. The liver sinusoids of *C3*^-/-^, *CRIg*/*CR3*^-/-^ and *Masp2*^-/-^ mice were imaged by intravital microscopy as in 2D. n = 2. **E.** Complement-dependent bacterial capture by KCs *in vitro*. Primary mouse KCs were incubated with *Spn-*19F in the presence of 10% serum for 30 min. *Spn-*14 was used as a negative control. KC-bound bacteria are presented as the percentage of total bacteria. MOI = 1; n = 3-6. **F.** The importance of the complement system in the protection against *Spn-*19F. Various complement-deficient mice were i.v. infected with 10^6^ CFU of *Spn-*19F and the survival is presented. n = 5. **G.** The model for the FCN-A-triggered complement-dependent and -independent pathways for bacterial capture by liver macrophages. FCN-A activates C3 on bacterial surface upon binding to the capsule through the lectin pathway. The representative data are presented as mean ± SD, and the statistical differences were determined by Two-way ANOVA with Tukey’s multiple comparisons test (A-C), multiple *t* tests (D and E) and log-rank test (F). **, *P* < 0.01; ****, *P* < 0.0001; ns, no significant difference.

We next determined how KCs capture C3-opsonized bacteria by performing infection with mice lacking CRIg and/or CR3, since CRIg and CR3 are the major complement receptors of KCs ^34, 49^. While *CRIg*^-/-^mice showed partial but significant impairment in hepatic clearance of *Spn-*19F, *CR3*^-/-^mice phenotypically behaved as WT mice (**Fig. 5B**, left panel), indicating CRIg as the major complement receptor on KCs that captures C3-opsonized bacteria. Further experiments revealed a minor role of CR3 in bacterial capture, since mice lacking both CR3 and CRIg (*CR3*^-/-^*/CRIg*^-/-^) displayed a more pronounced deficiency than *CRIg*^-/-^ mice. Infection with a high dose of *Spn-*19F also confirmed the importance of the complement receptors in mediating hepatic capture of C3-opsonized bacteria CRIg and CR3 (**Fig. 5B**, right panel).

Because human ficolins activate C3 via the lectin pathway *in vitro* ^45^, we assessed the role of this pathway in FCN-A-mediated bacterial clearance using mice lacking MASP1 or MASP2, two MBL-associated serine proteases that trigger the C3 activation upon the lectin binding to carbohydrate structures on microbial surface ^50^. Both *Masp1*^-/-^ and *Masp2*^-/-^ mice displayed a partial impairment in *Spn-*19F clearance at an infection dose of 10^6^ CFU (**Fig. 5C**, left panel), and the complete loss of such the activity when the dose was increased to 10^7^ CFU (right panel). These data indicated that ficolin-A activates C3 on bacterial surface via the lectin pathway.

The functional importance of the complement system in executing FCN-A-triggered bacterial capture in the liver was demonstrated by IVM imaging of mouse liver sinusoids. Bacterial immobilization onto KCs was dramatically reduced in *C3^-/-^*, *Masp2*^-/-^ and *CR3^-/-^/CRIg^-/-^* mice (**Fig. 5D**; **Video 9**). Under the *in vitro* conditions, primary mouse KCs abundantly captured *Spn-*19F bacteria in the presence of serum from WT mice, but not the serum from *C3^-/-^* or *Masp2*^-/-^ mice (**Fig. 5E**). The essential role of the complement system in FCN-A-mediated immunity was also manifested by the 100% mortality of *C3^-/-^* and *Masp2*^-/-^ mice post infection with a non-lethal dose of *Spn-*19F (**Figs. 5F**; **S4D**). By comparison, *CR3^-/-^CRIg^-/-^*mice fully survived the infection with 10^6^ CFU *Spn-*19F (**Fig. 5F**), which was consistent with milder phenotype of complement receptor-deficient mice in bacterial clearance (**Fig. 5B**, left panel). These mice showed a higher susceptibility to intravenous infection of 10^7^ CFU *Spn-*19F as compared with WT mice, with a 60% mortality (**Fig. S4E**). The functional linkage between ficolin-A and the complement system was further verified with serotype-9N pneumococci and serotype-f *H. influenzae* (**Figs. S4F-I**). These works revealed that FCN-A-binding to the capsule leads to C3 deposition on bacterial surface via the lectin pathway, and subsequent hepatic capture of C3-opsonized bacteria; FCN-A attachment to the capsule also engages an uncharacterized complement-independent mechanism of bacterial capture in the liver (**Fig. 5G**).

### Ficolin-A recognizes the *N*-acetylated monosaccharides of capsular polysaccharides

Since the fibrinogen-like domain of FCN-A is responsible for carbohydrate recognition (**Fig. 6A**, left panel) ^51^, we analyzed this segment to understand how the protein recognizes structurally diverse CPSs. Consistent with the current knowledge that ficolins recognize acetylated monosaccharides ^29, 30, 52^, we noticed that the repeating units of all FCN-A-binding CPSs contain at least one *N-*acetylated monosaccharides (**Fig. 6A**, right panel) ^32^. rFCN-A bound to the CPSs of the nine FCN-A-sensitive serotypes, whereas it did not show significant binding to CPS11A, which lacks an *N*-acetyl group (**Fig. 6B**). The ELISA data also showed significant variations in the level of FCN-A binding to different capsules, suggesting that ficolin-A possess variable levels of physical affinity to different capsules. In particular, CPS9N exhibited the lowest binding to FCN-A. Competitive binding assays using monosaccharides to inhibit FCN-A and CPS19F interaction demonstrated that the *N*-acetylhexosamine serves as the recognition motif for ficolin-A (**Fig. 6C**). These results suggested that ficolin-A binds to structurally diverse capsules by recognizing *N-*acetylated monosaccharides of CPSs.

**Figure 6.**
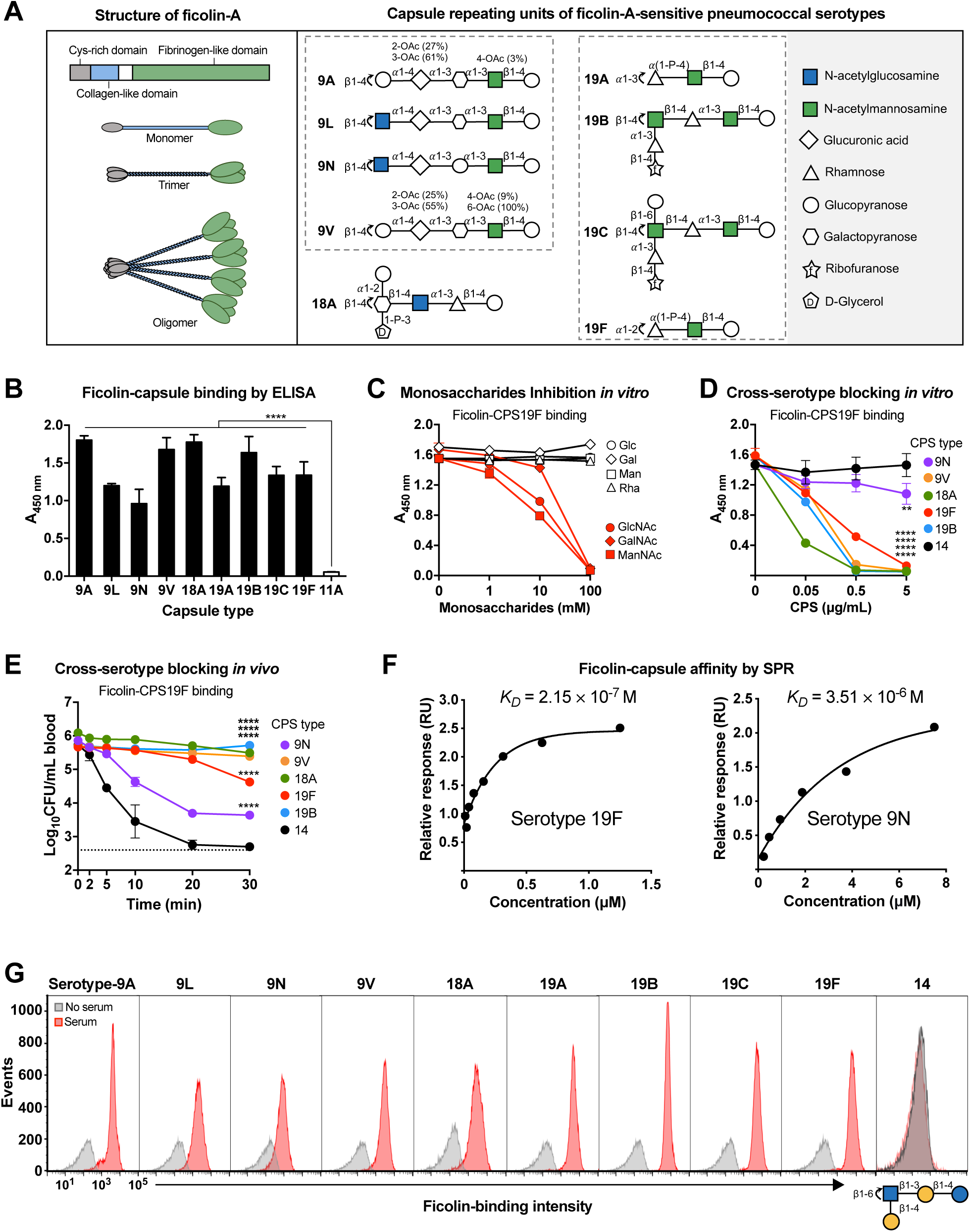
Specific recognition of the *N*-acetylated capsules by mouse ficolin-A. **A.** The predicted structures of ficolin-A (left) and repeat units of FCN-A-recognizable capsules (right). **B.** rFCN-A binding to free pneumococcal CPSs was assessed by ELISA. n = 3. **C.** Competitive blocking of rFCN-A and CPS19F binding by monosaccharides were determined by ELISA. n = 3. **D.** Inhibition of rFCN-A-CPS19F binding by free CPSs was determined by preincubating with rFCN-A before being added to CPS19F-coated wells for ELISA. Serotype 14 was used as a negative control. n = 3. **E.** Blocking of the *Spn-*19F clearance from the bloodstream was determined by i.v. inoculation of 400 μg individual CPS 2 min before i.v. infection with 10^6^ CFU of *Spn-*19F to assess bacteremia kinetics. Dash line indicates the detection limit. n = 3. **F.** The affinities of rFCN-A to CPSs of serotypes 9N and 19F determined by surface plasma resonance (SPR), are presented as dissociate constants (*K*_D_). n = 2. **G.** Differential binding of FCN-A to *N-*acetylated pneumococcal surfaces were assessed by flow cytometry. The repeat unit of serotype 14, which is not bound by FCN-A, is indicated at the bottom. The representative data are presented as mean ± SD, and the statistical differences were determined by Two-way ANOVA with Tukey’s multiple comparisons test (E and F). **, *P* < 0.01; ****, *P* < 0.0001; ns, no significant difference.

To define if FCN-A binds to different capsules with the same single interface, we used free CPSs of selected serotypes to block the FCN-A-CPS19F interaction. As compared with CPS of FCN-A-insensitive serotype-14, the counterparts of serotype-9N, -9V, -18A and -19B significantly blocked the binding of rFCN-A to CPS19F in a dose-dependent manner, although CPS9V showed a much weaker inhibition (**Fig. 6D**). This cross-serotype inhibition was confirmed *in vivo*. Intravenous inoculation with CPSs of the FCN-A-sensitive serotypes before infection significantly impaired the clearance of *Spn-*19F from the circulation (**Fig. 6E**). As a negative control, similar treatment with purified CPS14 did not show obvious blocking effect. These experiments strongly suggested that ficolin-A binds to structurally diverse capsules with the same ligand-binding interface.

The significant differences in the binding levels of FCN-A to various CPSs (Fig. 6B) and in the inhibitory effects of CPSs on FCN-A binding (Fig. 6D) suggest that the binding affinities between the CPSs and FCN-A may vary. To quantify the biochemical interactions of FCN-A and *N-*acetylated capsules, we determined the binding affinity of FCN-A to CPS19F and CPS9N (with the lowest binding level in Fig. 6B) using surface plasmon resonance (SPR). The SPR experiment results revealed an affinity or dissociation constant (*K_D_*) of 2.15 × 10^-7^ M for CPS19F (**Fig. 6F**). Consistent with the ELISA result (Figs. 6B and 6D), rFCN-A showed a relatively binding to CSP9N (*K_D_* = 3.51 × 10^-6^ M), which is 15.3-fold higher than the counterpart for CPS19F. This result showed that FCN-A binds to capsules with variable levels of affinity.

To characterize the capsule-binding spectrum of FCN-A, we performed deposition of rFCN-A on the surface of 30 pneumococcal serotypes with *N*-acetylated CPSs in our collection by flow cytometry, which representing 52.6% of all the 57 pneumococcal serotypes with *N*-acetylated CPSs ^32, 53^ (**Table S3**). While rFCN-A abundantly deposited to all the 9 FCN-A-sensitive serotypes, the other 21 *N*-acetylated serotypes did not display obvious FCN-A binding (4, 5, 7B, 7C, 10B, 10C, 10F, 11B, 11C, 11D, 12A, 12F, 13, 14, 15A, 15F, 29, 33B, 33C, 33D and 47A) (**Figs. 6G** and **S5**). The FCN-A-sensitive serotypes each contains one (9A, 9V, 18A, 19A and19F) or two (9N, 9L, 19B and 19C) *N*-acetylated CPS hexosamines, which include *N*-acetylglucosamine (9L, 9N and 18A) and/or *N*-acetylmannosamine groups (all of the 9 serotypes except for 18A) (**Fig. S6A**). Additional experiment also showed that FCN-A bound to the surface of type f *H. influenzae* (**Fig. S5**), whose capsule contains two *N*-acetylgalactosamine residues ^54^ (**Fig. S6B**). This result indicated that mouse ficolin-A recognizes the *N*-acetylated glucosamine, mannosamine and galactosamine CPS components of the FCN-A-sensitive serotypes. However, these *N*-acetylhexoamines along are not sufficient for the FCN-A recognition since these groups also exist in the CPSs of the 21 serotypes lacking the FCN-A binding activity. While some of these FCN-A-insensitive capsules contain more than two *N*-acetylhexosamines in the repeat units (types 4, 5, 12A and 12F), many of them possess a single *N*-acetylated hexosamine that is attached to a branch monosaccharide (types 7C, 10B, 10C, 10F, 11B, 11C, 14 and 47A), surrounded by adjacent sugars with anomeric configuration (α/β) (types 7B, 15A and 15F), or linked to a furanose (types 13, 29, 33B, 33C and 33D) (**Fig. S5**). These data indicated that FCN-A recognizes the *N*-acetylated hexosamines in the context of local chemical conditions.

### Ficolin-2 enables human Kupffer cells to capture pneumococci with *O-*acetylated capsules

Human ficolin-2 (FCN-2) is functional ortholog of mouse FCN-A, based on their commonalities in sequence, hepatic production and blood localization ^10^. FCN-2 is reported to bind to *O-*acetylated CPSs of 14 pneumococcal serotypes (**Fig. S7** and **Table S3**) ^18–20^. In light of the FCN-A-triggered bacterial clearance, we characterized potential interaction of FCN-2 with the FCN-A-sensitive serotypes of *S. pneumoniae*. Consistent with the previous observation ^18–20^, flow cytometry showed serotype-specific deposition to *O-*acetylated serotype-11A, -33A, -35A and -42 pneumococci, but not to 12 *N*-acetylated serotypes, including nine FCN-A-sensitive serotypes and three FCN-A-insensitive serotypes (12A, 14, 29) (**Figs. 7A** and **S8A**). In an opposite manner, mouse FCN-A did not bind the ten FCN-2-binding serotypes (11A, 11D, 15F, 31, 33A, 35A, 35C, 42, 47A and 47F) (**Table S3**), which agrees with the normal clearance of the FCN-2-binding serotypes in FCN-A-deficient mice (**Table S2**). The results were further verified by ELISA (**Fig. S8B**). These lines of evidence have uncovered that the human and mouse ficolin orthologs recognize distinct structural patterns of capsules. While human ficolin-2 targets the *O-*acetylated monosaccharides, mouse ficolin-A binds to *N-*acetylated sugars.

**Figure 7.**
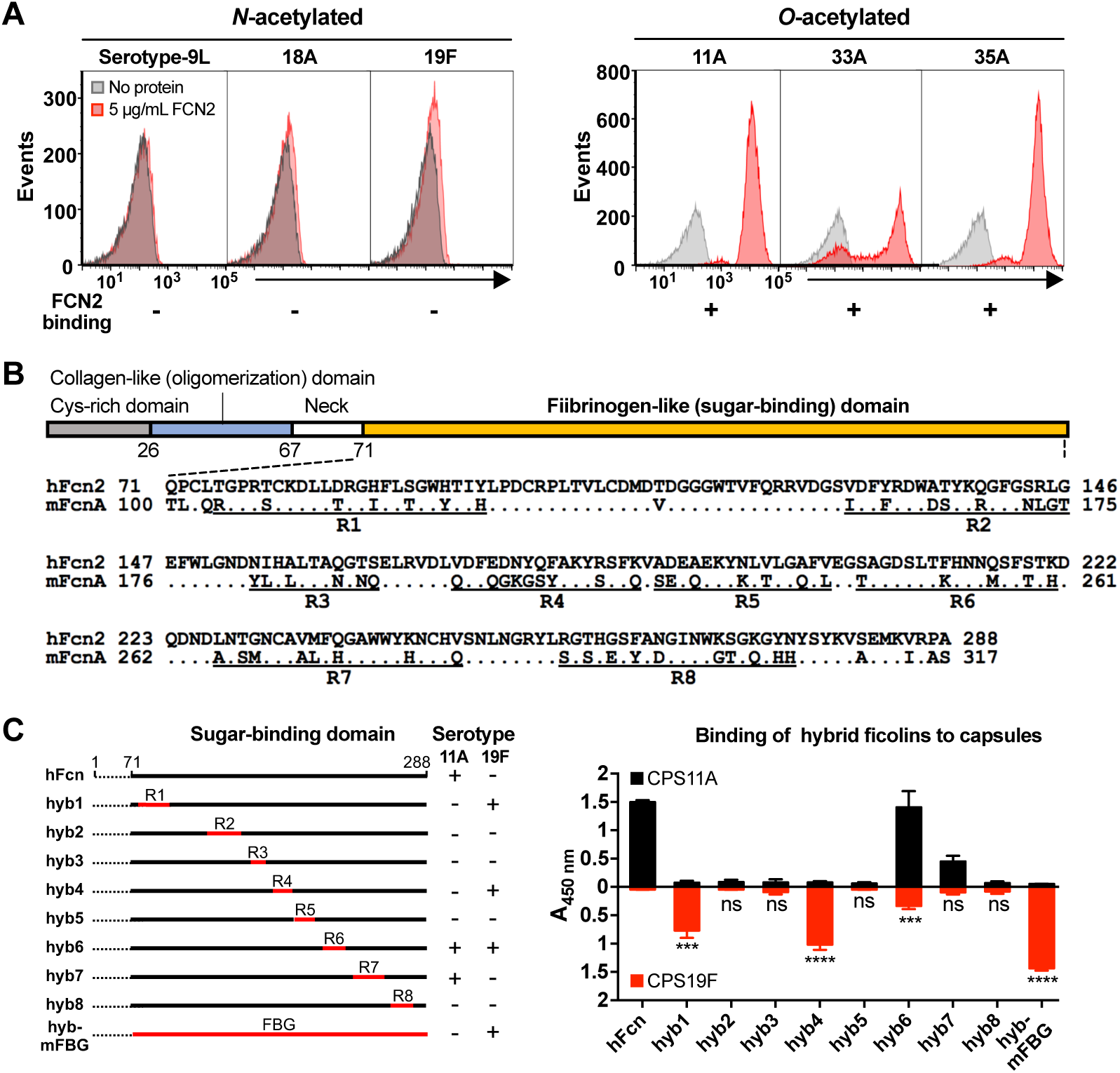
Specific recognition of the *O-*acetylated capsules by human ficolin-2. **A.** Capsule acetylation mode-dependent deposition of human ficolin-2 to pneumococcal surface is indicated as positive (+) or negative (-) binding at the bottom of each serotype. **B.** Sequence alignment between the sugar-binding domains of human ficolin-2 (hFcn) and mouse ficolin-A (mFcn). The divergent sequences were partitioned into eight regions labeled R1-R8. **C.** The effect of single-region substitutions between hFcn and mFcn on capsular recognition specificity was assessed by ELISA (right). The construction of ficolin variants is in the context of FCN-2 and the diagram are shown (left). The representative data are presented as mean ± SD, and the statistical differences were determined by multiple *t* tests (C). ***, *P* < 0.001; ****, *P* < 0.0001; ns, no significant difference.

Since the ligand specificities of the ficolins are defined by their fibrinogen-like domains ^51^, we characterized the functional motifs that contribute the distinct ligand specificities of the human and mouse ficolins. The sequence diversity between the two proteins is concentrated in eight regions (R1-R8 in **Fig. 7B**), based on the secondary structural elements of the human FCN-2 ^55^. We constructed eight chimeric FCN-2 proteins (hyb1-8) by individually replacing the eight divergent regions with the corresponding sequences of FCN-A (**Fig. 7C**, left panel). Except for hyb6 that retained the binding activity of CPS11A (the ligand FCN-2), all of the chimeras virtually failed to bind to the capsule (**Figs. 7C**, right panel). However, hyb1 and hyb4 showed significant binding to CPS19F, the ligand of FCN-A, thus switching the ligand specificity from CPS11A to CPS19F. The remaining six chimeras showed no (hyb2, hy3, hyb5, hyb7 and hyb8) or marginal (hyb6) CPS19F-binding activity. This result indicated that regions R1, R4 and R6 contribute to the substrate-binding specificity of FCN-2.

## DISCUSSION

Ficolins bind to a wide range of bacteria, fungi and viruses, and activate the complement system under the *in vitro* conditions ^14^. These glycan-binding proteins are therefore regarded as pathogen-recognition receptors. However, it remains unclear if the lectin activities of ficolins contribute to the microbial capture and clearance in the process of infections. This work has uncovered that plasma ficolins of human (ficolin-2) and mouse (ficolin-A) enable the liver macrophages to rapidly capture blood-borne encapsulated bacteria by recognizing capsular polysaccharides. Binding of mouse ficolin-A to the *N-*acetylated capsules drives swift trapping and killing of encapsulated bacteria by KCs in the liver sinusoids. Ficolin-A-deficient mice became hyper-susceptible to septic infections of the targetable capsule types. To the best of our knowledge, this study presents the first *in vivo* evidence that ficolins directly drives phagocytic capture of pathogens during the infection process.

### Ficolin-A confers a potent anti-bacterial innate immunity

Based on the broad binding of ficolins to microbial glycans, a large body of literature has been generated to validate the hypothesis that these lectins possess anti-infection functions ^14^. This notion is supported by the *in vitro* activation of the complement system upon glycan binding of human ficolin-1, 2 and 3 ^16, 28, 29, 41, 56^. Other *in vit*ro studies have linked ficolin binding to opsonophagocytosis of encapsulated bacteria ^19, 23, 46, 57^. By using ficolin-A-deficient mice, this work has shown that the natural blood level of ficolin-A confers a sterilizing immunity against septic infections by a wide range of encapsulated bacteria. As an example, ficolin-A deficiency made mice 13,570-fold more susceptible to *Sp*n-19F infection. This immunity level is equivalent to the protection induced by pneumococcal polysaccharide vaccine^58.^

The potency of the ficolin-mediated immunity differs among the capsule types. Taken the pneumococci as examples, the clearance of serotype-9A, -9V, -19A and -19F was completely abolished in FCN-A knockout mice, but only partial defect was observed with serotype-18A and-19B. This trend was also obvious in bacterial hepatic capture/killing and host survival. This variability indicates that ficolin-A is the sole receptor for some capsule types, but the other types are redundantly recognized by additional receptor(s). In addition, the capsule binding affinity of ficolin-A apparently contributed to the serotype-specific potency of the protein.

### Ficolins cover a broad range of pathogens

Our limited screening has identified 12 ficolin-A-binding capsules of Gram-positive and -negative bacteria (9 types of *S. pneumoniae,* 2 types of *H. influenzae* and 1 type of *E. coli*). FCN-A-deficient mice showed a significant impairment in the hepatic clearance of these serotypes. While the precise spectrum of the ficolin-mediated immunity remains to be defined, we believe that FCN-A should be able to recognize additional capsule types beyond those discovered in this work. Ficolin-A has been shown to bind to certain fungal pathogens ^59^. Likewise, the human ortholog (ficolin-2) is reported to recognize the capsules of *S. pneumoniae*^18–20^, *S. agalactiae* ^23–25^, and *S. aureus* ^18^. Moreover, ficolin-2 also interacts with *Salmonella* Typhimurium ^60^, *Mycobacterium avium* ^61^, *Aspergillus fumigatus* ^62^ and hepatitis C virus ^63^, although the microbial ligands for these interactions were not characterized. Finally, the previous studies have also documented *in vitro* binding of other human and mouse ficolins to pathogenic microorganisms ^14^.

This broad pathogen coverage of ficolins is reminiscent of the wide immunity of plasma natural antibodies and CRP. Natural antibodies activate the hepatic capture of enteropathogenic *E. coli* in mice ^64^. Likewise, natural antibodies trigger rapid clearance of serotype-10 and -39 *S. pneumoniae* and serotype-K50 *K. pneumoniae* by mouse KCs via recognizing the β1-6-linked galactose branch in the capsular polysaccharides ^36^. CRP recognizes the capsules of 20 bacteria, and thereby enables the capsule type-specific capture of these pathogens in the liver sinusoids of mice ^37^. Our recent study has also revealed that capsular polysaccharide-induced antibodies also achieve protective immunity by enabling KCs to capture encapsulated bacteria in the liver sinusoids, but these antibodies are highly specific to the individual capsule types represented by the vaccines ^58^. In this sense, the wide pathogen coverage is a major advantage of the innate capsule receptors.

### Ficolin-A engages liver macrophages by multiple pathways

In line with the functional relationship between ficolins and the complement system ^45^, we found that the hepatic clearance of ficolin-A-bound bacteria involves the activation of the complement lectin pathway. This conclusion is based on the significant defect of C3- or MASP-deficient mice in the hepatic clearance of FCN-A-sensitive pneumococci. Along the same line, mice lacking CRIg and CR3, the major complement receptors on KCs ^34, 49^, showed a similar degree of deficiency in this activity. The combined data indicate that the activated C3 serves as the functional bridge to physically link the ficolin-bound bacteria and complement receptors on KCs. However, we’ve also noticed a partial clearance of ficolin-bound bacteria in the livers of mice lacking C3, MASP1/2 or complement receptors at a relatively low inoculum. This observation shows that ficolin-A is able to shuffle bacteria to Kupffer cells via an uncharacterized complement-independent mode. This immuno-redundancy has also been reported for natural antibody and CRP-mediated bacterial clearance in the liver ^36, 37, 64^. This redundancy appears to have an evolutionary advantage in maximizing the anti-bacterial power of pathogen pattern recognition receptors.

### The glycan-binding domains of ficolins shape the host specificity of encapsulated bacteria

It has been reported that mouse ficolin-A and human ficolin-2 bind to a similar spectrum to bacteria and fungi ^26^. However, this study has revealed that these functional ficolin orthologs recognize a completely different sets of capsules. All of the 11 ficolin-A-sensitive capsules with the known repeating unit structures contain *N-*acetylated monosaccharides (e.g. ManNAc, GlcNAc and GalNAc), but only four of them carry *O-*acetylation (e.g. serotype-19A and -19V *S. pneumonia*e and serotype-c and -f *H. influenzae*). In contrast, ficolin-A did not bind to any of the 10 tested *O-*acetylated pneumococcal capsules that are recognized by human ficolin-2 ^18–20^. Accordingly, all of the 10 ficolin-2-recognizable pneumococcal serotypes tested in this work were normally cleared in FCNA-deficient mice. In an opposite manner, recombinant ficolin-2 only bound to the capsules with only *O-*acetylated monosaccharides, but not to any of the *N-*acetylated counterparts that are recognized by ficolin-A. This finding suggests that mouse ficolin-A selectively binds to *N-*acetylated capsules while the human counterpart favors *O*-acetylated ones. The structural basis for this strong ligand selectivity of these ficolin orthologs remains to be further defined.

### Plasma capsule receptors are essential for maintaining the blood sterility

The sterilizing effect of mouse ficolin-A has a strong implication in the molecular basis of blood sterility. The liver is known as a major part of the reticuloendothelial system to trap and clear blood-borne bacteria and other foreign particles ^65–70^. At the cellular level, the liver resident macrophages – KCs are found to be the dominant phagocytes in capturing bloodstream microbes ^71, 72^. The dominant expression of several receptors shapes the unique capacity of KCs in microbial capture, including the asialoglycoprotein receptor ^34^ and CRIg complement receptor ^37, 49, 58, 73–77^. However, it is largely unclear how Kupffer cells target such a large range of microbes with the limited receptor repertoire. In context of the recently discovered plasma capsule receptors (e.g. CRP and natural antibodies) ^36, 37^, this work has uncovered the immune power and microbial coverage of a single plasma protein in driving hepatic clearance of invading bacteria from the blood circulation. A common feature of these plasma capsule receptors is that they all utilize the complement system as a functional linkage to the complement receptors on KCs. This modular arrangement maximizes the usage of the limited receptor repertoire of KCs. It is certain that these capsule receptors only represent the tip of the receptor iceberg in maintaining the blood sterility, but the emerging conceptual framework will aid the discovery of new sterilizing receptors and the comprehensive understanding of the immunological basis for maintaining blood sterility.

### The perspectives on the anti-bacterial immunity of human ficolins

Ficolins are relatively new comers in the field of immunology since the first ficolin was identified in 1993 ^9^. Based on their recognition of microbial glycans, human ficolins have been expensively investigated *in vitro*, in terms of activating the complement protein C3 on microbes and promoting bacterial opsonophagocytosis. However, human ficolins have not been demonstrated to promote bacterial clearance *in vivo*. Based on the potent anti-bacterial immunity of ficolin-A in mouse sepsis model, we also observed that human ficolin-2 enhances serotype-specific bacterial capture by primary human liver macrophages. These data strongly suggest that human ficolin-2 principally fulfills a similar sterilizing immunity as mouse ficolin-A. This notion is strongly supported by the current literature. First, human ficolin-2 recognizes the capsules of many pneumococcal serotypes and other bacteria ^15,18–20, 22–25^; ficolin-2 binding to capsules leads to the activation of the lectin complement pathway and phagocytic uptake of *S. pneumoniae* ^19, 20, 45, 46^ and *S. agalactiae* ^78^. Second, human epidemiology studies have associated serum ficolin levels with the serotype-specific susceptibility to encapsulated bacteria. The ficolin-2-recognizable serotype-11A *S. pneumoniae* is among the serotypes with the lowest invasiveness in childhood invasive pneumococcal disease (IPD), but is more prevalent in IPD cases of older adults with relatively lower levels of serum ficolin-2 ^19, 20^. Likewise, the low ficolin-2 level in umbilical cord serum is associated with preterm delivery and low birth weight ^79^.

### Limitations of the study

A major limitation is that all the *in vivo* infection works were conducted in mouse sepsis model; the resulting information may not be directly extrapolated into the functions of human ficolins. However, both human ficolin-2 and mouse ficolin-A showed a broad recognition of many capsule types. Moreover, ficolin-2 promoted *in vitro* serotype-specific capture of *S. pneumoniae* by primary human Kupffer cells. In this context, we anticipate that ficolin-2 also constitutes a sterilizing immunity in the human bloodstream. It should be noted that human ficolin-2 recognizes a completely different set of pathogens from mouse ficolin-A. Furthermore, human has an additional plasma ficolin (ficolin-3). While ficolin-3 has not been documented to recognize any bacterial capsules, it forms heterocomplexes with ficolin-2 ^80^, which may impact the biology of ficolin-2. These lines of information indicate that plasma ficolins of human and animals fulfill similar anti-bacterial functions, but the molecular details of their actions may vary.

## MATERIALS AND METHODS

### Bacterial strains and cultivation

All bacteria used in this study are summarized in Tables S4. *S. pneumoniae* was cultured in Todd-Hewitt broth (THB) with 0.5% yeast extract (THY) or tryptic soy agar (TSA) plates containing 4% sheep blood as described ^81^. *H. influenzae* was grown in brain-heart infusion (BHI) broth or BHI agar with 10 μg/mL hemin (Sigma) and 10 μg/mL nicotinamide adenine dinucleotide (NAD) (Sigma)as described ^82^. *E. coli* was cultivated in Luria-Bertani (LB) broth or on LB agar plates as described ^35^. For DH5α derivative strains, when required, the following were added: kanamycin (50 μg/mL) or ampicillin (100 μg/mL). Prior to use, all bacterial strains were aliquoted in broth medium containing 25% glycerol (OD₆₂₀ = 0.5-0.6) and stored at -80°C.

### Mutagenesis

The pneumococcal capsule-switch strains were constructed by natural transformation as described ^34^. Briefly, the type 19F strain TH2740, which was rendered streptomycin-resistant by transformation with the rpsL1 allele, was used as the recipient background strain. The capsule biosynthesis genes in the *cps* locus were replaced by the Janus cassette (JC) via XbaI/XhoI digestion and ligation of the JC amplicons with the *cps* up- and downstream sequence amplicons, generating the kanamycin-resistant Δ*cps*::JC background strain TH7338. For capsule switching, the target *cps* locus was amplified from the genomic DNA of donor strains using PrimeSTAR GXL DNA polymerase (Takara). The resulting amplicon was subsequently introduced into the Δ*cps*::JC background strain to generate the corresponding capsule-switched strains. The specific primers and procedures are listed in Tables S5 and S6, respectively.

### Purification of capsular polysaccharide (CPS)

Bacterial CPSs were purified from broth cultures using a zwitterionic detergent elution method and quantified by phenol-sulfuric acid method as described ^34, 35^.

### Isolation of liver non-parenchymal cells (NPCs)

Liver NPCs were isolated as described ^34^. In brief, the liver of euthanized mice was perfused from the portal vein with digestion buffer (HBSS with 0.5 mg/mL collagenase IV, 20 μg/mL DNase I, and 0.5 mM CaCl_2_). The liver was minced into small pieces (< 2 mm) and incubated in digestion buffer at 37°C for 30 min. After digestion, the liver homogenate was filtered through a 70-μm strainer and washed. The residual red blood cells were lysed by resuspending the pellet in RBC lysis solution (BioLegend) and incubating it on ice for 1 min. The hepatocytes were removed by centrifugation at 50 g for 2 min and the liver NPCs in the supernatant were collected.

### Screening for CPS19F-binding proteins

Pneumococcal CPS19F-binding proteins were screened by an affinity pull-down strategy using CPS19F-coated beads as described ^83^. Briefly, the membrane proteins were extracted from the liver NPCs using Mem-PER™ Plus Membrane Protein Extraction Kit (Thermo Scientific) following the manufacturer’s instructions. Purified polysaccharides were conjugated to carboxyl beads. To enrich CPS-binding proteins, 10^8^ CPS-coated beads were mixed with 100 μg membrane proteins in 500 μL phosphate buffered saline with Tween-20 (PBST) supplemented with 20% serum and 2 mM CaCl_2_ and MgCl_2_. The mixtures were rotated at room temperature (RT) for 1 hour (hr). CPS8-coated beads were used as a negative control since CPS8 cannot be recognized by KCs. Proteins bound on CPS-beads were subjected to quantitative protein identification by mass spectrometry. Protein abundance was compared between CPS19F- and CPS8-coated beads, and proteins enriched at least 2-fold by CPS19F-beads were considered as CPS19F-binding candidates. Proteins identified by mass spectrometry following affinity screening with CPS19F and CPS8 were listed in Table S7.

### Production of recombinant ficolins

Mouse ficolin-A (rFCN-A) and human ficolin-2 (rFCN-2) were expressed as Strep-tagged recombinant proteins in HEK293F cells principally as described ^37^. Briefly, the full-length coding sequence of FCN-A was amplified from the murine liver cDNA using primers Pr20048/Pr20049 and cloned into the NotI/BamHI site of the pCMV-chikv-strepII vector ^84^ in *E. coli* DH5α, yielding plasmid pFCN-2 in strain TH17167. *E. coli* containing pFCN-2 was grown in LB containing 50 μg/mL kanamycin. Recombinant protein was produced by transient transfection of HEK293F cells with pFCN-2, and purified from the culture supernatant using Strep-Tactin sepharose resin (IBA), and eluted with 2.5 mM desthiobiotin in TBS-Ca²⁺ buffer (100 mM Tris, 150 mM NaCl, 2 mM CaCl₂, pH 8.0). The target protein-containing fractions were pooled and buffered exchanged using a 3 kDa MWCO ultrafiltration device to remove desthiobiotin. Protein concentration was determined using the BCA Assay Kit (Solarbio). rFCN-2 was generated in a similar manner. The coding sequence of rFCN-2 was synthesized using the accession NM_004108 (Beijing Xianghong Biotechnology), and used to construct protein-expressing plasmid pFCN-2 in strain TH17168.

Chimeric proteins with exchanged FBDs between human FCN-2 and mouse FCN-A (hyb-hFBD and hyb-mFBD) were constructed using plasmid pcDNA3.4 as vector. The primers and templates are illustrated in Fig. S9A. These amplicons were assembled using the ClonExpress Ultra One Step Cloning Kit (Vazyme), which were transformed into *E. col* and selected on LB agar in the presence of 100 μg/mL ampicillin.

Chimeric FCN-2 proteins containing FCN-A segments (hyb1-8) were constructed using pFCN-2 as templates. Sequences from regions 1 to 8 of the *Fcna* gene were incorporated into the 5’ ends of the primers to linearize the plasmid by PCR amplification. The primers and templates are listed in Fig. S9B. The amplicons were phosphorylated by T4 polynucleotide kinase, ligated by T4 DNA ligase, and transformed into *E. coli* for resistance to 50 μg/mL kanamycin. These chimeric ficolins were expressed and purified as describe above. The primers and plasmids used in this study are described in Tables S5 and S8, respectively.

### Enzyme-linked immunosorbent assay (ELISA)

The interaction between ficolins and CPS was assessed by ELISA principally as described ^37^. Briefly, CPSs (10 μg/mL in PBS) were coated on the 96-well plates and blocked with 5% non-fat milk in TBST-Ca^2+^ buffer (TBS- Ca^2+^ with 0.1% Tween-20). The plates were incubated sequentially with 100 µL of Strep-tagged recombinant ficolins, mouse anti-Strep-tag monoclonal antibody (Huaxing Bio, 1:2000), and horseradish peroxidase (HRP)-conjugated goat anti-mouse IgG (Huaxing Bio, 1:2000), with extensive washing after each step using 200 µL TBST. Finally, ficolin-CPS binding was detected by adding 100 μL of TMB peroxidase substrate (TIANGEN), followed by the addition of 100 μL of 1 M phosphoric acid to stop the reaction, and measuring the optical density at 450 nm with a microplate reader (BioTek). All the reagents were diluted in TBST-Ca^2+^ buffer. For competitive inhibition, free CPSs or monosaccharides were incubated with Strep-tagged recombinant ficolins at RT for 1 hr before adding to CPS-coated wells.

To determine the complement activation by ficolin-capsule interactions, CPS-coated microtiter wells were filled with 100 μL of 20% mouse serum in TBST-Ca^2+^ buffer. The C3 deposition on CPSs was detected using a 1:2000 dilution of HRP-conjugated goat IgG antibody against mouse complement C3 (MP Biomedicals) after incubation at 37℃ for various durations.

### Detection of *in vitro* bacterial binding by ficolins

The ficolin-A binding to the intact pneumococci was analyzed by fluorescence microscopy as described ^36^. Briefly, bacteria (5 × 10^7^ CFU) in 100 μL PBS were stained with 10 μg of AF647-NHS ester at RT for 30 min. After washing, the bacteria were sequentially incubated with 50% (v/v) mouse serum in PBS containing 2 mM CaCl₂ at 37°C for 30 min, a rabbit anti-ficolin-A antibody (Cloud-Clone Corp, 1:500) for 15 min, and an FITC-conjugated goat anti-rabbit IgG (Easybio, 1:500) for 15 min in the dark at RT, with extensive washing between the steps. Bacterial fluorescence was visualized using an Olympus SpinSR10 confocal microscope with two laser excitation wavelengths (488 nm and 640 nm).

The interaction between ficolins and bacteria was also assessed by flow cytometry as described^37^. Briefly, bacteria (2 × 10^6^ CFU) were incubated with 10% normal serum or 50 µg/mL recombinant ficolins A in Ringer’s solution for 15 min at RT. Serum FCN-A deposition was detected with a rabbit anti-ficolin-A polyclonal antibody (Cloud-Clone Corp, 1:100) and a FITC-conjugated goat anti-rabbit IgG (1:100); rFCN-A binding with a mouse anti-Strep-tag monoclonal antibody (Huaxing Bio, 1:1000) and a goat anti-mouse IgG (1:1000). Bacterial fluorescence was analyzed using a FongCyte flow cytometer (Challenbio).

### Mouse infection

All the infection experiments were conducted in C57BL/6 or CD1 mice (6-8 weeks old) from Vital River according to the animal protocol approved by the Institutional Animal Care and Use Committee of Tsinghua University (Protocol No. 22-ZJR1). All gene-deficient mice were maintained on the C57BL/6 background. *Fcna*^-/-^ and *Fcnb*^-/-^ mice were kindly provided by Dr. Xulong Zhang in the Capital Medical University ^85^. C3-deficient (C3^-/-^) and B cell-deficient (μMT) mice were obtained from the Jackson Laboratory (Bar Harbor, Maine, USA). CR3-deficient (CR3^-/-^ or *Itgam*^-/-^) mice were generated by CRISPR/Cas9 system as described ^58^.CRIg -deficient (CRIg^-/-^ or *Vsig4*^-/-^) mice originated from Genetech (USA) ^49^. CR3 and CRIg double deficient (CR3/CRIg^-/-^) mouse lineages were generated by cross breeding of CR3^-/-^ and CRIg^-/-^ mice. MASP1-deficient (MASP1^-/-^) and MASP2-deficient (MASP2^-/-^) mice were purchased from Jiangsu GemPharmatech (Suzhou, China) and Model Organisms Center (Shanghai, China), respectively.

Septic infections were carried out by i.v. infections as described ^34^. Briefly, mice were i.v. injected with 10^6^ CFU of bacteria in 100 μL of Ringer solution. Bacterial early clearance from the bloodstream was assessed by retroorbital bleeding at various time points post infection. Organ bacterial burden was determined by enumerating CFUs of homogenized organs. Total bacterial load in each mouse was calculated as the sum of CFU values in the blood and major organs. Bacterial 50% clearance time (CT_50_) was calculated by nonlinear regression analysis of bacteremia kinetics using the formula T = ln{(1-50/Plateau)/(-K)}, in which Plateau and K were generated by one-phase association of the bacterial clearance ration using GraphPad Prism. Survival rate of mice was monitored for 7 days post infection.

### Intravital microscopy (IVM)

IVM imaging of mouse liver sinusoids was performed as described ^34^. In brief, lLSECs and KCs were stained by i.v. injection of AF594-conjugated anti-CD31 antibody (Biolegend) and AF647-conjugated anti-F4/80 antibody (Invitrogen), respectively, before i.v. administration of FITC-labeled bacteria. Images of the liver vasculatures were acquired with Leica TCS-SP8 confocal microscope using 10×/0.45 NA and 20×/0.80 NA HC PL APO objectives 10-15 min post infection.

Photomultiplier tubes (PMTs) and hybrid photo detectors (HyD) were used to detect fluorescence signals (600 × 600 pixels for time-lapse series and 1024 × 1024 pixels for photographs). At least 5-10 ten random fields of view (FOV) were captured to calculate bacterial number per FOV. Leica Biosystems software was used for image and movie processing (LAS X Life Science).

### Bacterial binding to primary Kupffer cells

Bacterial binding to primary mouse and human Kupffer cells was assessed as described ^34^. Briefly, liver NPCs were resuspended in serum-free RPMI 1640 medium (Corning) and seeded into 96-well plates at 5 × 10^4^ cells/well in 100 μL. KCs were enriched by cellular adherence to the plastic surface after incubation at 37°C, 5% CO_2_ for 30 min; nonadherent cells removed by gentle washing with RPMI 1640 medium. KCs were co-cultured with bacteria at MOI 1:1 in 100 µL of RPMI 1640 medium in the presence or absence of 10% normal serum or 10 µg/mL of recombinant ficolins. After incubation at 37°C for 30 min, supernatant was collected to quantify CFU of free bacteria. KCs were lysed with 100 µL of ice-cold H₂O at 4°C for 15 min to release and quantify the cell-associated bacteria. The binding ratio is defined as the percentage of cell-associated CFUs out of the total CFUs.

### Surface plasmon resonance (SPR)

The binding affinity between rFCN-A and CPSs was analyzed by SPR using a Biacore 8K+ instrument as described with minor modifications ^86^.CM5 Briefly, the sensor chip was first activated with a mixture of 1-Ethyl-3-(3-dimethylaminopropyl) carbodiimide (EDC) and *N*-Hydroxysuccinimide (NHS). rFCN-A (50 µg/mL in 10 mM sodium acetate, pH 4.0) was immobilized onto the chip surface via amine coupling. Remaining active groups were blocked with ethanolamine-HCl. All binding steps were performed using TBST-Ca^2+^ buffer as the running buffer at 25°C.

Serially diluted polysaccharide solutions at empirically determined concentrations were injected as analytes at a flow rate of 30 µL/min. The contact time was 120 seconds (s); the dissociation time 120 s. The sensor surface was regenerated with a 30-second pulse of 500 mM NaCl at a flow rate of 30 µL/min between cycles. The equilibrium responses at each polysaccharide concentration were plotted against analyte concentration; the equilibrium dissociation constant (*K*_D_) determined by fitting the binding isotherm to a steady-state model using the Biacore Insight Evaluation software.

### Sequence analysis

DNA and amino acid sequences were analyzed using DNASTAR Lasergene (version 18.0.3.2) for Macintosh.

### Protein structural prediction

The structures of FCN-A, hyb1 and hyb4 were predicted by AlphaFold3 ^87^; the structural figures were prepared using UCSF ChimeraX ^88^.

### Data analysis

Statistical analyses were performed using GraphPad Prism software version 8.0.0, and all data were expressed as Mean ± SEM unless otherwise stated. The unpaired Student’s t-test and multiple t-test was used for comparing two and multiple groups, respectively. Data involving two independent variables were analyzed by two-way ANOVA followed by Tukey’s multiple comparison test. Mouse survival was analyzed by log-rank (Mantel-Cox) test. *P* value < 0.05 was considered as significant (not significant, ns; \**P* < 0.05; \*\**P* < 0.01; \*\*\**P* < 0.001; \*\*\*\**P* < 0.0001).

## ACKNOWLEDGMENTS

We are grateful to Drs. Kai-Hu Yao, Yingchun Xu, Jie Feng, Fen Qu and Lijun Wang for providing bacterial strains, and Xulong Zhang for providing *Fcna*^-/-^ and *Fcnb*^-/-^ mice. We thank the Tsinghua research platforms for assistance in animal experimentation (Laboratory Animal Research Center), flow cytometry and IVM imaging (Center for Cell Biology), and SPR experiments and protein mass spectrometry (Technology Center for Protein Sciences).

## FUNDING

This work was supported by the following grants: National Key Research and Development Program of China grant 2023YFc2306301 (J-RZ), National Natural Science Foundation of China 32300153 (XH), 82330071 (J-RZ) and 82522055 (HA), Tsinghua University Initiative Scientific Research Program 20243080033 (J-RZ) and Conducting Surveys, Research and Risk Assessment on Zoonotic Diseases 0733-24095348 (J-RZ).

## COMPETING INTERESTS

There are no competing interests with this publication.

## DATA AND MATERIALS AVAILABILITY

All data and materials used in this study will be made available under the conditions of material transfer agreements.

## AUTHOR CONTRIBUTIONS

Conceptualization: XH, JF, HA, J-RZ; methodology: XH, JF, HA, JH, YL, YX; investigation: XH, HC, JF, KL; materials: L-TS, RZ, PY, HT, ST; visualization: XH, HC, JF, KL; funding acquisition: J-RZ, XH; project administration: XH, J-RZ; supervision: XH, J-RZ; writing: XH, JF, YX, J-RZ.

---

**Figure S1.**
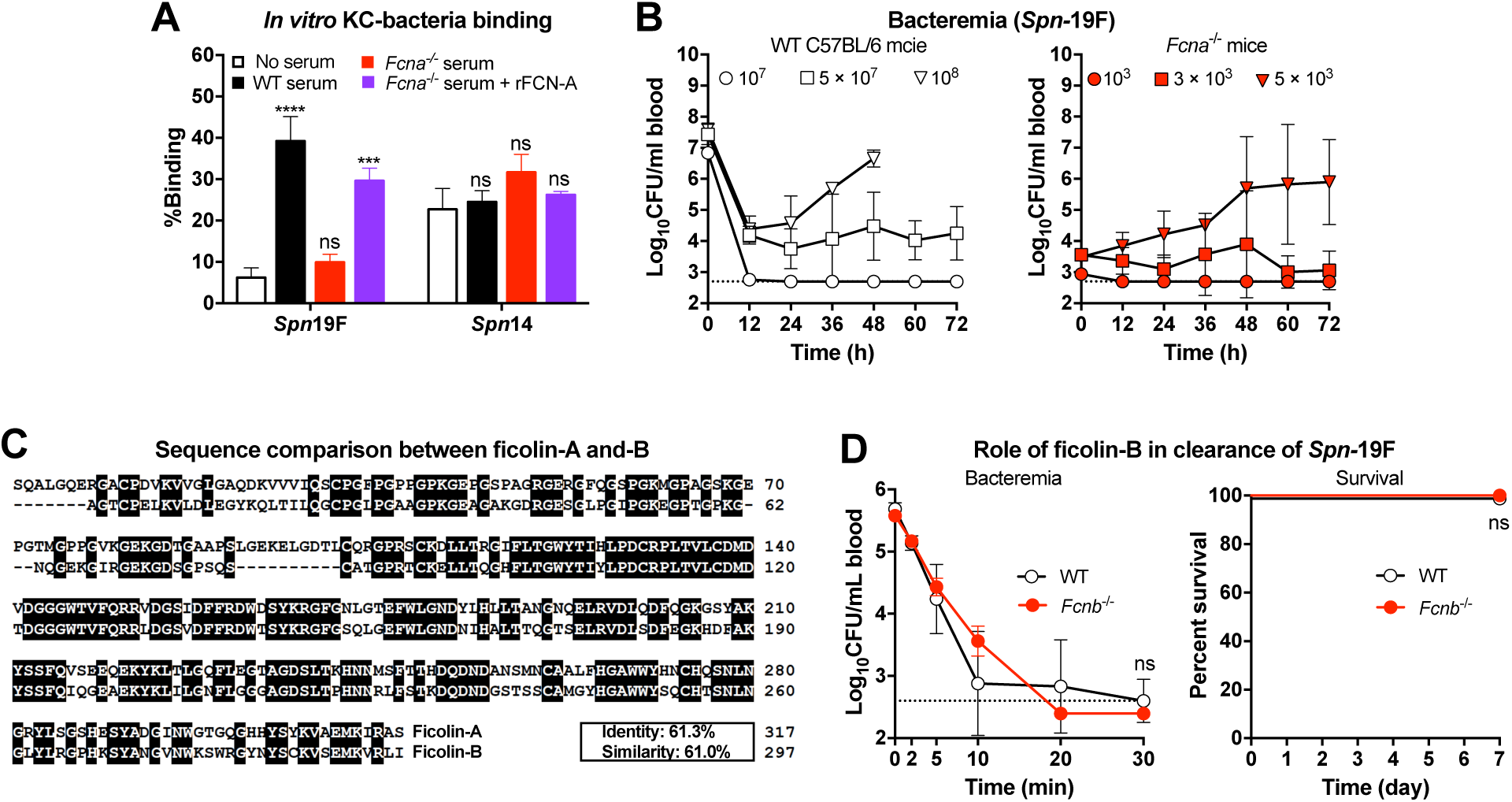
Role of mouse ficolin in defense against *Spn-*19F infection (related to Fig. 2). **A.** FCNA-mediated primary mouse KC binding to *Spn-*19F were determined *in vitro* in the presence of 10% serum alone or in combination with 20 μg/mL rFCN-A. Serum-free medium was used as a negative control, and *Spn-*14 was used as a positive control, which binds directly to KC without serum. The percentage of binding was presented as the ratio of KC-associated CFU to total CFU per well after 30 min incubation. MOI = 1; n = 3-6. **B.** Bacterial load in the blood of WT (left) and *Fcna*^-/-^ (right) mice i.v. infected with various dose of *Spn-*19F was monitored. Dotted line, detection limit; n = 5. **C.** Sequence alignment of murine ficolin-A and ficolin-B. **D.** The role of ficolin-B in bacterial clearance and host defense against *Spn-*19F infection was determined by plating blood CFU of mice at various time points post i.v. infection with 10^6^ CFU of *Spn-*19F (left) and monitoring survival of mice for 7 days (right). n = 5. The representative data are presented as mean ± SD, and the statistical differences were determined by multiple *t* test (A). \*\*\**P* < 0.001; \*\*\*\**P* < 0.0001; ns, no significant difference.

**Figure S2.**
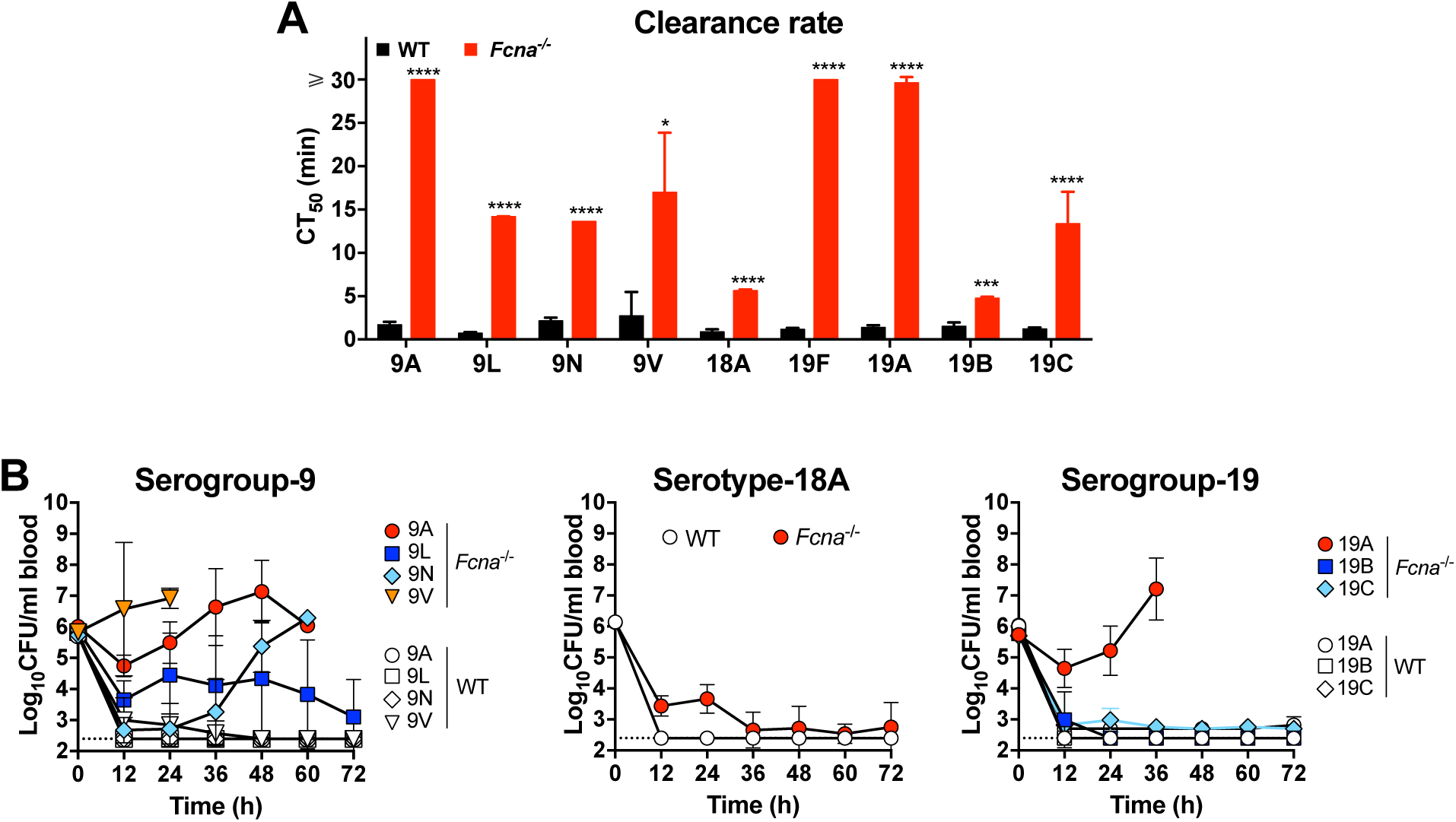
Ficolin-A-mediated host resistance to diverse pneumococcal serotypes (related to Fig. 3). **A.** The clearance rates of pneumococci from the bloodstream of mice i.v. infected with 10^6^ CFU pneumococci were presented as the time to clear 50% of inoculum from the bloodstream. n = 3. **B-D**. The role of Ficolin-A in bacteremia kinetics following infection with serogroup-9, serotype-18A, or serogroup-19 in mice was determined by plating CFU in blood of mice at intervals of 12 h post infection as in (A). Dash line indicates the detection limit. n = 5. The representative data are presented as mean ± SD, and the statistical differences were determined by multiple *t* test (A). \**P* < 0.5; \*\*\**P* < 0.001; \*\*\*\**P* < 0.0001.

**Figure S3.**
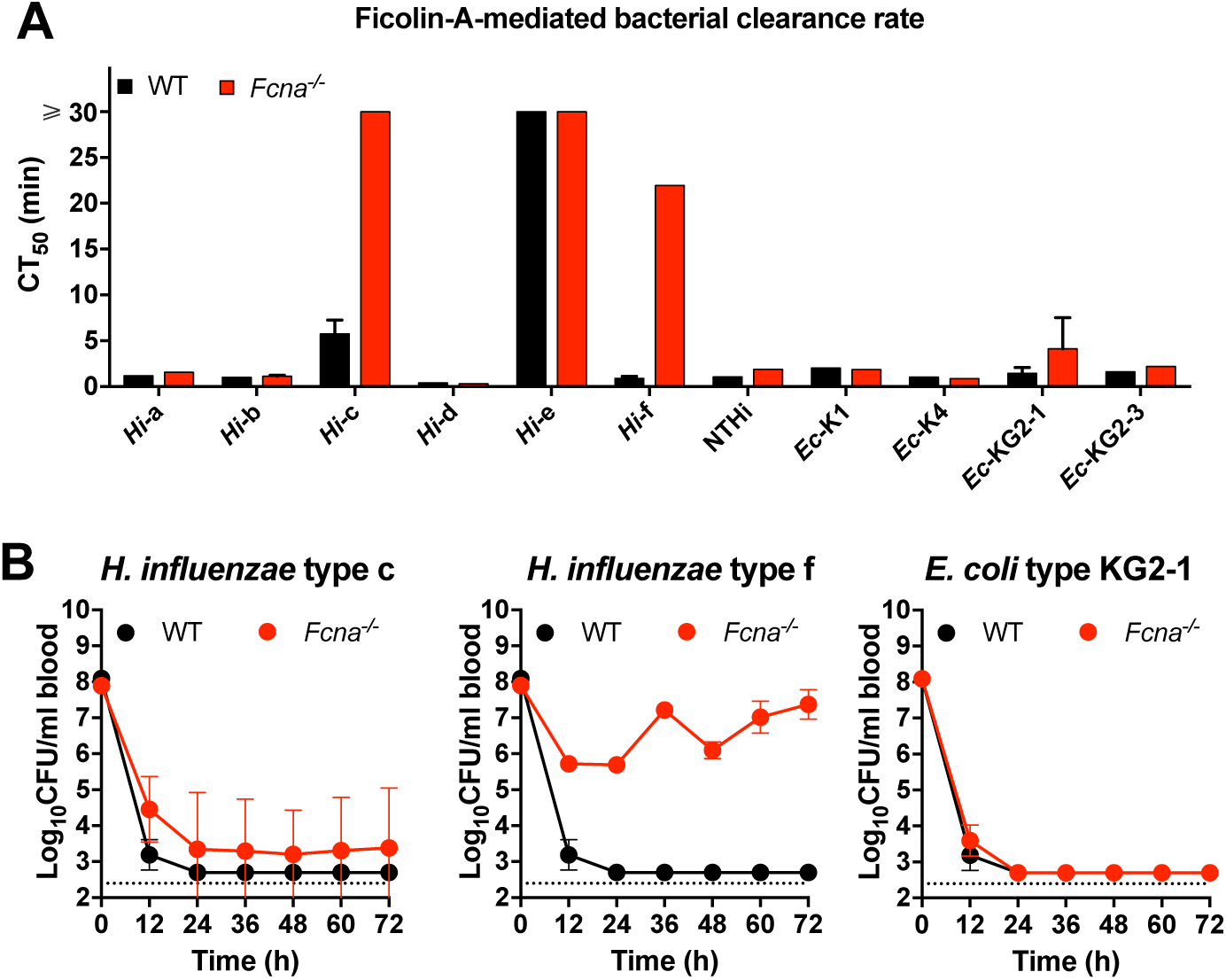
Importance of ficolin-A in host resistance to Gram-negative bacteria (related to Fig. 4). **A.** 50% clearance time of Gram-negative bacteria from the blood in mice with 10^6^ CFU i.v. infection was calculated as in Fig. 4A. n = 1-3 **B.** Bacteremia kinetics of mice i.v. inoculated with 10^8^ CFU bacteria was monitored at 12-hour intervals post infection. Dash line indicates the detection limit. n = 5. The representative data are presented as mean ± SD.

**Figure S4.**
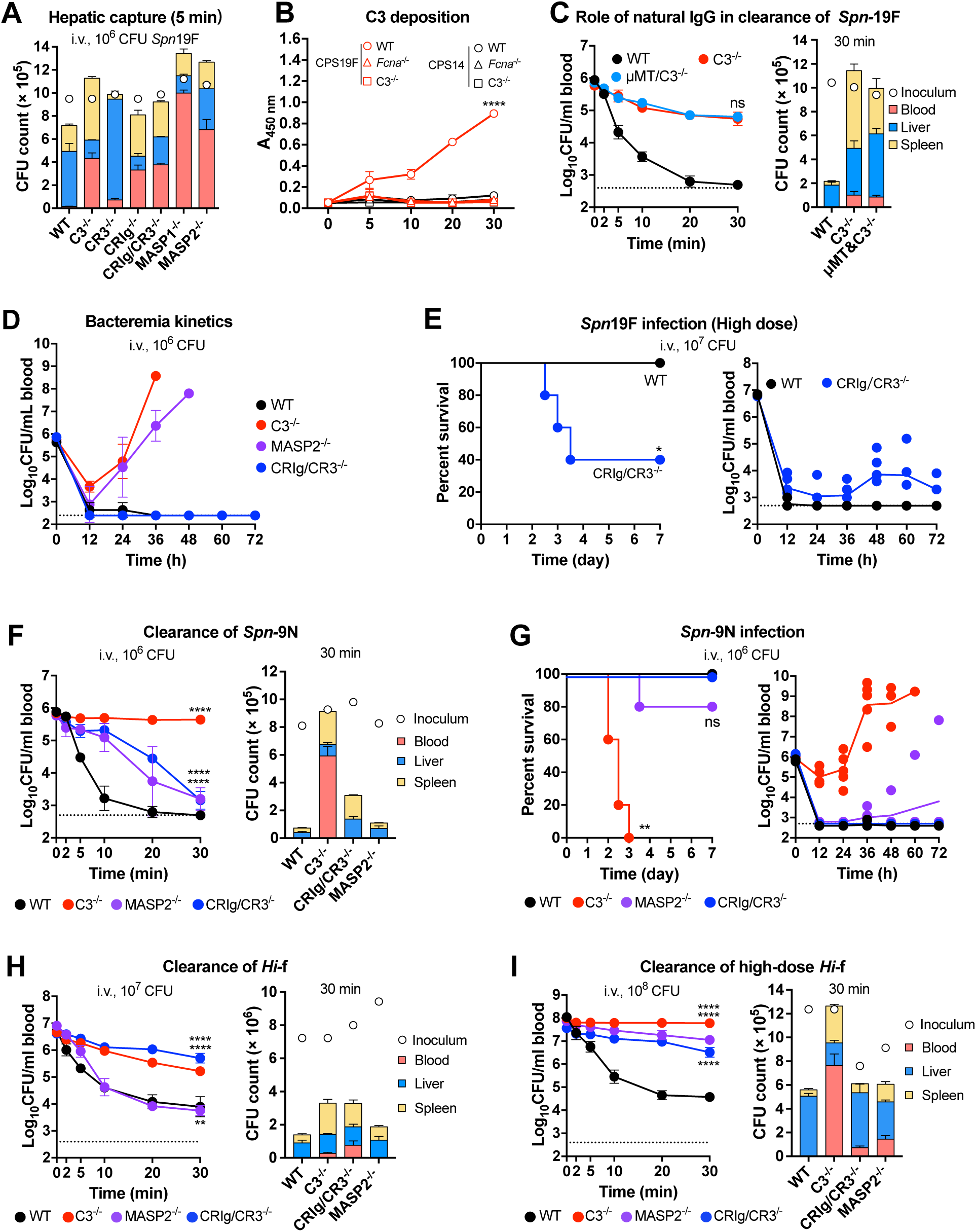
Role of the complement lectin pathway to ficolin-A-activated immunity against bacterial infection (related to Fig. 5). **A.** Bacterial distribution in blood, liver and spleen of C3, MASP1/2 or complement receptor-deficient mice i.v. infected with 10^6^ CFU of *Spn-*19F were determined separately at 5 min post infection, to demonstrate the role of complement system in hepatic capture of *Spn-*19F. n = 3. **B.** C3 deposition on CPS19F and CPS8 were detected after incubation at various time points in the presence of 10% serum from WT, *Fcna*^-/-^ or *C3*^-/-^ mice, respectively. n = 3. **C.** Bacteremia kinetics (left) and bacterial distribution (right) of *C3*^-/-^ and natural antibody-deficient *C3*^-/-^ mice (μMT&*C3*^-/-^) infected as in (B) were determined to investigate the role of natural antibody in clearance of FCN-A-opsonized *Spn-*19F by the C3-independent pathway. Dash line indicates the detection limit. n = 3. **D.** Long-term bacteremia kinetics of WT, *C3*^-/-^, *Masp2*^-/-^ and *CRIg*/*CR3*^-/-^mice were monitored by plating and counting the bacteria CFU in blood at various time points after infection. n = 5. **E.** Survival rates (left) and blood bacteria (right) of *CRIg*^-/-^*CR3*^-/-^ mice infected with high dose (10^7^ CFU) of *Spn-*19F were determined. n = 5. **F.** Bacteremia kinetics (left) and bacterial distribution (right) of complement component-deficient mice (*C3*^-/-^, *Masp2*^-/-^ and *CRIg*/*CR3*^-/-^) were determined to indicate the role of complement system in FCN-A-mediated clearance of *Spn-*9N. n = 3. **G.** Survival rates (left) and long-term bacteremia kinetics (right) of complement component-deficient mice infected with 10^6^ CFU of *Spn-*9N were determined. n = 5. **H**, **I**. The role of complement system in clearance of 10^7^ (**H**) and 10^8^ (**I**) CFU of *Hi-*f was determined as in (F). n =3-5.

**Figure S5.**
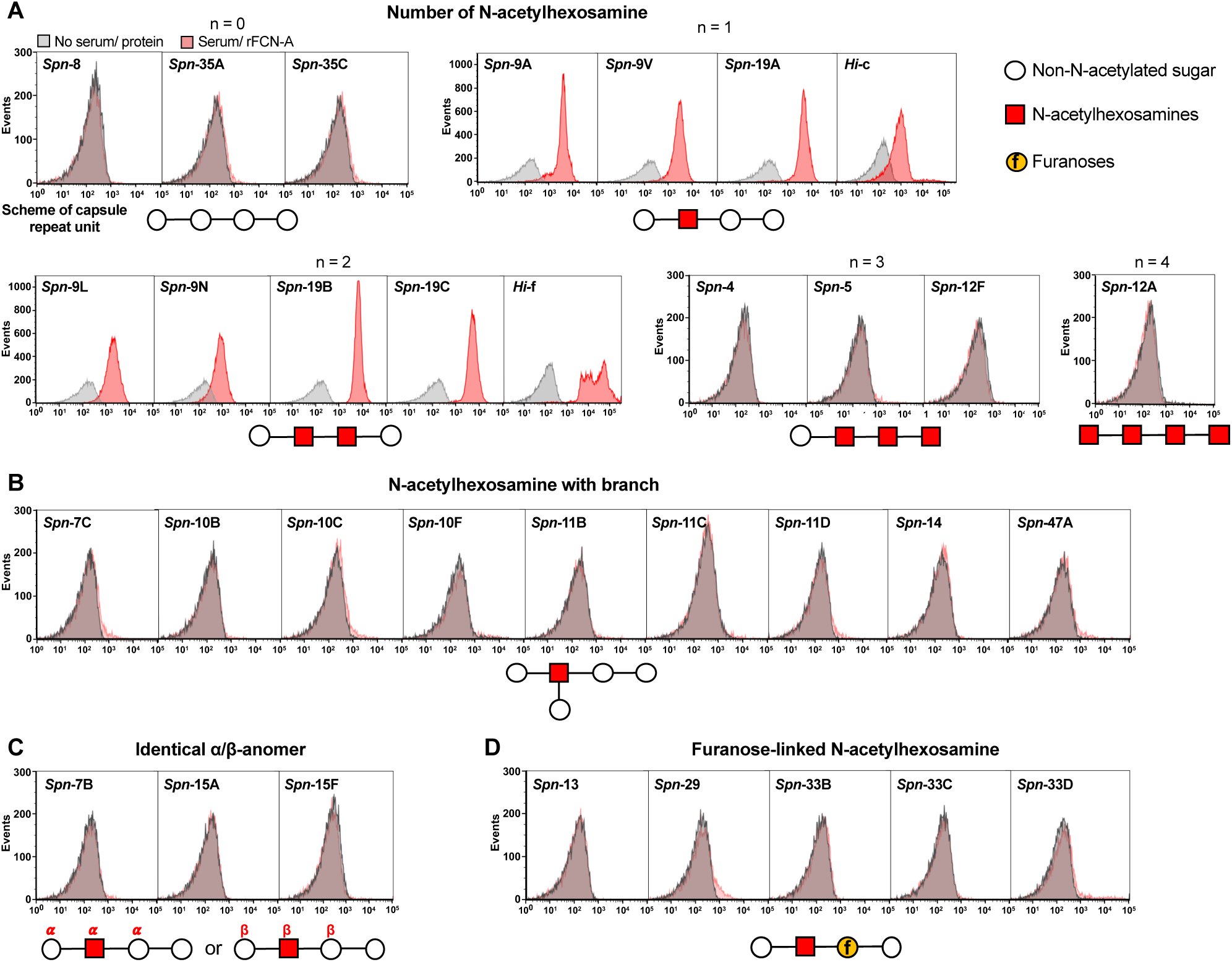
Specific binding of ficolin-A to pneumococci carrying various forms of *N*-acetyl CPSs (related to Fig. 6). Binding of murine FCN-A to pneumococci was quantified by flow cytometry based on distinct structural features of the N-acetylhexosamine residue in capsule repeat unit: number of residues (**A**), branch pattern (**B**), anomeric configuration (α/β) (**C**), and structural features of adjacent sugars (**D**). Schematics of the capsular repeat units are shown below each panel.

**Figure S6.**
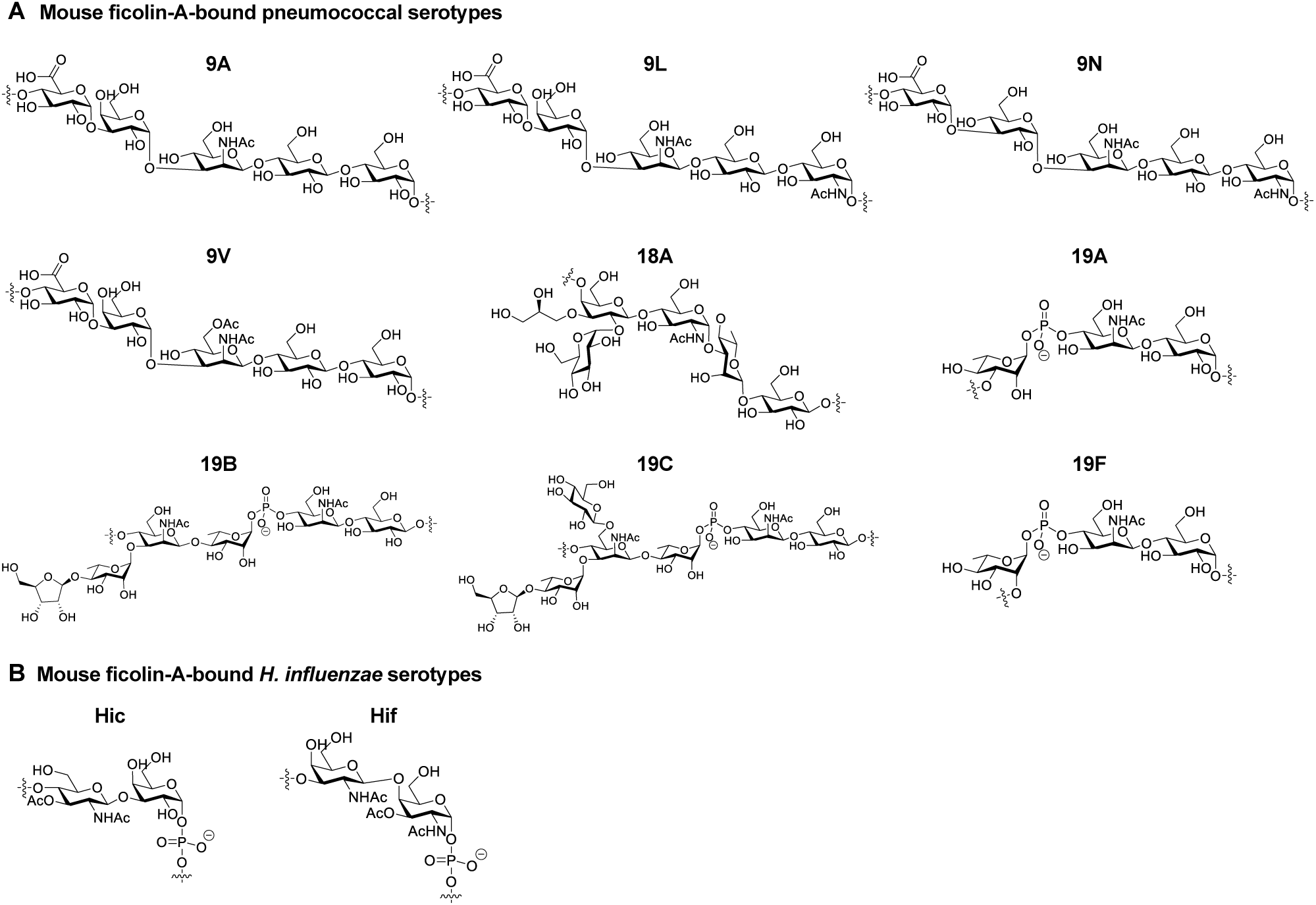
Chemical structures of ficolin-A-binding capsules (related to Fig. 6). A. Chemical structures of FCN-A-binding pneumococcal capsules (serogroup-9, serotype-18A and serogroup-19) B. Chemical structures of FCN-A-binding *H. influenzae* types c and f capsules.

**Figure S7.**
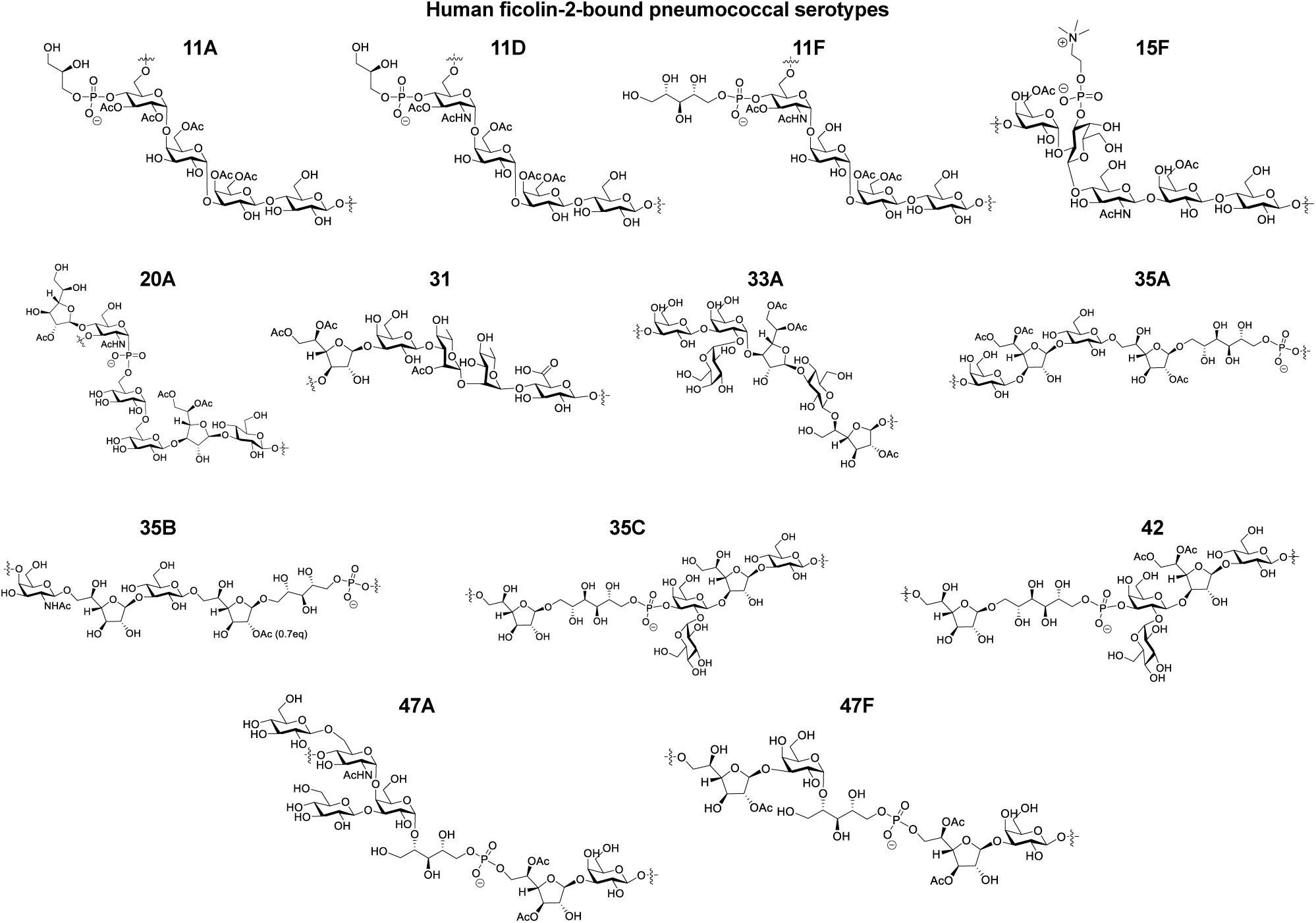
Chemical structures of human ficolin-2-binding capsules (related to Fig. 7).

**Figure S8.**
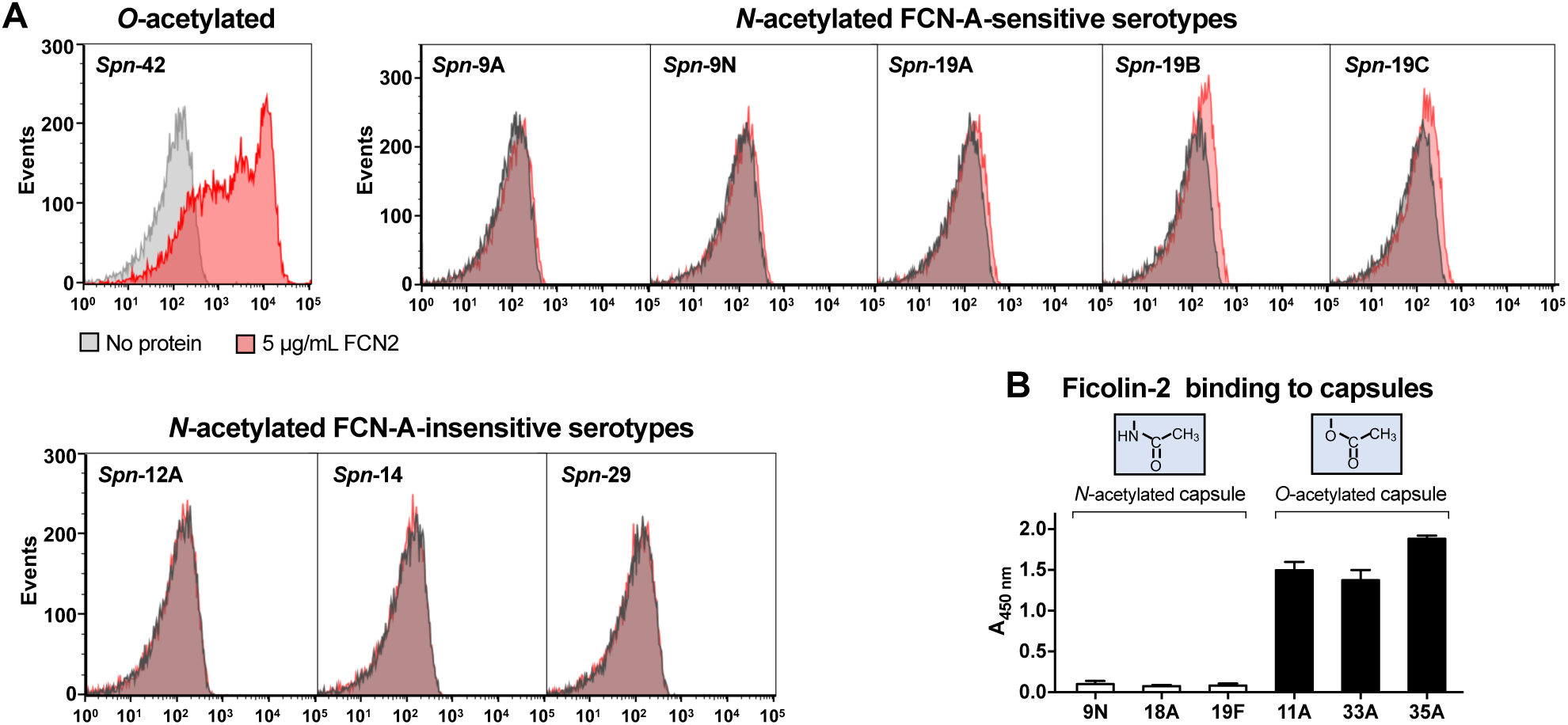
Specific recognition of the *O*-acetylated pneumococci by human ficolin-2 (related to Fig. 7). **A.** Binding of rFCN-2 to *O*- or *N*-acetylated pneumococci was quantified by flow cytometry. **B.** rFCN-2 binding to free pneumococcal CPSs was assessed by ELISA. n = 3.

**Figure S9.**
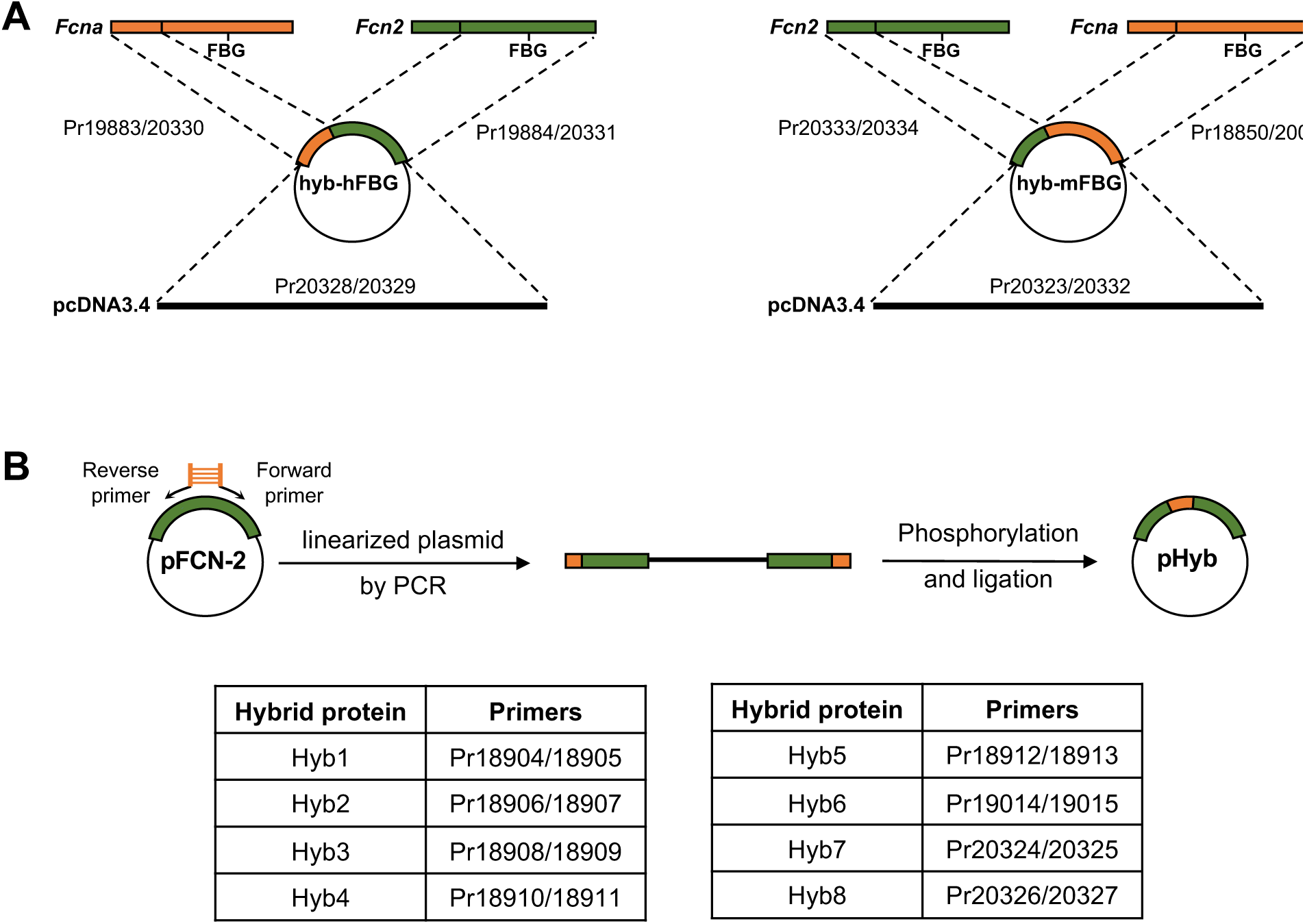
Schematic diagram of experimental procedure for construction of hybrid ficolins (related to Fig. 7). **A.** Construction of hybrid proteins hyb-hFBG (left) and hyb-mFBG (right). Plasmids phyb-hFBG and phyb-mFBG encode hybrid proteins consisting of the *N*-terminus of one ficolin and the fibrinogen-like (FBG) domain of the other. **B.** Construction of hybrid proteins Hyb1 to Hyb 8. The sequences of regions 1 to 8 were incorporated into the 5’ ends of the corresponding primers, respectively, and then integrated into the plasmid via phosphorylation and ligation of the linearized plasmid.

## SUPPLEMENTAL INFORMATION LIST

### **<u>Supplemental figures</u>** (included in the main text)

Figure S1. Role of mouse ficolin in defense against *Spn*-19F infection (related to Fig. 2)

Figure S2. Ficolin A-mediated host resistance to diverse pneumococcal serotypes (related to Fig. 3)

Figure S3. Importance of ficolin-A in host resistance to Gram-negative bacteria (related to Fig. 4)

Figure S4. Role of the complement lectin pathway to ficolin A-activated immunity against bacterial infection (related to Fig. 5)

Figure S5. Specific binding of ficolin-A to pneumococci carrying various forms of *N-*acetyl CPSs (related to Fig. 6)

Figure S6. Chemical structures of ficolin-A-binding capsules (related to Fig. 6)

Figure S7. Chemical structures of human ficolin-2-binding capsules (related to Fig. 7)

Figure S8. Specific recognition of the *O-*acetyl pneumococci by human ficolin-2 (related to Fig. 7)

Figure S9. Schematic diagram of experimental procedure for construction of hybrid ficolins (related to Fig. 7)

### **<u>Supplemental tables</u>** (included in the supplemental information file)

Table S1. CPS19F-enriched mouse proteins identified by mass spectrometry **Table S2**. Early clearance characteristics of bacteria in the bloodstream of mice **Table S3**. Deposition of ficolin-A and ficolin-2 on the pneumococci

Table S4. Bacterial strains and derivatives used in this study

Table S5. Primers used in this study

Table S6. Information for constructions of strains used in this study

Table S7. Proteins identified by mass spectrometry following affinity screening with CPS19F and CPS8

Table S8. Plasmids used in this study.

### **<u>Supplemental videos</u>** (submitted as separate files)

**Video 1**. Impaired FCN-A-mediated capture of serotype 19F pneumococci in liver sinusoids of *Fcna*^-/-^ mice.

**Video 2**. Effective capture of serotype-14 pneumococci in liver sinusoids of *Fcna*^-/-^ mice. **Video 3**. Impaired FCN-A-mediated capture of serotype-9N pneumococci in liver sinusoids of *Fcna*^-/-^ mice.

**Video 4**. Impaired FCN-A-mediated capture of serotype-18A pneumococci in liver sinusoids of *Fcna*^-/-^ mice.

**Video 5**. Impaired FCN-A-mediated capture of serotype-19A pneumococci in liver sinusoids of *Fcna*^-/-^ mice.

**Video 6**. Impaired FCN-A-mediated capture of serotype c *H. influenza*e in liver sinusoids of *Fcna*^-/-^ mice.

**Video 7**. Impaired FCN-A-mediated capture of serotype f *H. influenza*e in liver sinusoids of *Fcna*^-/-^ mice.

**Video 8**. Impaired FCN-A-mediated capture of serotype KG2-1 *E. coli* in liver sinusoids of *Fcna*^-/-^ mice.

**Video 9**. Impaired FCN-A-mediated capture of *Spn-*19F in the liver sinusoids of *C3*^-/-^, *Masp2*^-/-^or CR3^-/-^/CRIg^-/-^mice

## Notes

### Competing Interest Statement

The authors have declared no competing interest.

