## Supplementary material for "Plasma Ficolins Enable Liver Macrophages to Capture Blood-Borne Bacteria by Recognizing Capsular Polysaccharides": Table S4

**Table S4. Bacterial strains and derivatives used in this study**

| **Strain** | **Description** | | **Reference or source** |
| --- | --- | --- | --- |
| D39 | *Streptococcus pneumoniae* isolates, serotype 2 | | (An et al., 2022) |
| A66 | *S. pneumoniae* isolates, serotype 3 | | (An et al., 2022) |
| TH869 | *S. pneumoniae* isolates, serotype 38 | | (An et al., 2022) |
| TH870 | *S. pneumoniae* isolates, serotype 6A | | (An et al., 2022) |
| TH872 | *S. pneumoniae* isolates, serotype 22F | | (An et al., 2022) |
| TH882 | *S. pneumoniae* isolates, serotype 18C | | (An et al., 2022) |
| TH2734 | *S. pneumoniae* isolates, serotype 17F | | (An et al., 2022) |
| TH2737 | *S. pneumoniae* isolates, serotype 24A | | (An et al., 2022) |
| TH2740 | *S. pneumoniae* isolates, serotype 19F | | (An et al., 2022) |
| TH2918 | *S. pneumoniae* isolates, serotype 19A | | (An et al., 2022) |
| TH2932 | *S. pneumoniae* isolates, serotype 12F | | (An et al., 2022) |
| TH10833 | *S. pneumoniae* isolates, serotype 5 | | (An et al., 2022) |
| TH12927 | *S. pneumoniae* isolates, serotype 15A | | (An et al., 2022) |
| TH12932 | *S. pneumoniae* isolates, serotype 11A | | (An et al., 2022) |
| TH15984 | *S. pneumoniae* isolates, serotype 9N | | (An et al., 2022) |
| TH16764 | *S. pneumoniae* isolates, serotype 7B | China (Institute of Microbiology Chinese Academy of Sciences) | |
| TH16767 | *S. pneumoniae* isolates, serotype 7C | China (Institute of Microbiology Chinese Academy of Sciences) | |
| TH16770 | *S. pneumoniae* isolates, serotype 10B | China (Institute of Microbiology Chinese Academy of Sciences) | |
| TH16775 | *S. pneumoniae* isolates, serotype 11B | China (Institute of Microbiology Chinese Academy of Sciences) | |
| TH16777 | *S. pneumoniae* isolates, serotype 11C | China (Institute of Microbiology Chinese Academy of Sciences) | |
| TH16778 | *S. pneumoniae* isolates, serotype 13 | China (Institute of Microbiology Chinese Academy of Sciences) | |
| TH16781 | *S. pneumoniae* isolates, serotype 15F | China (Institute of Microbiology Chinese Academy of Sciences) | |
| TH16784 | *S. pneumoniae* isolates, serotype 16F | China (Institute of Microbiology Chinese Academy of Sciences) | |
| TH16788 | *S. pneumoniae* isolates, serotype 17A | China (Institute of Microbiology Chinese Academy of Sciences) | |
| TH16789 | *S. pneumoniae* isolates, serotype 18A | China (Institute of Microbiology Chinese Academy of Sciences) | |

**Table S4. Bacterial strains and derivatives used in this study (Continued)**

| **Strain** | **Description** | **Reference or source** | |
| --- | --- | --- | --- |
| TH16791 | *S. pneumoniae* isolates, serotype 18F | China (Institute of Microbiology Chinese Academy of Sciences) | |
| TH16792 | *S. pneumoniae* isolates, serotype 21 | China (Institute of Microbiology Chinese Academy of Sciences) | |
| TH16794 | *S. pneumoniae* isolates, serotype 22A | China (Institute of Microbiology Chinese Academy of Sciences) | |
| TH16801 | *S. pneumoniae* isolates, serotype 28F | China (Institute of Microbiology Chinese Academy of Sciences) | |
| TH16802 | *S. pneumoniae* isolates, serotype 29 | China (Institute of Microbiology Chinese Academy of Sciences) | |
| TH16804 | *S. pneumoniae* isolates, serotype 31 | China (Institute of Microbiology Chinese Academy of Sciences) | |
| TH16807 | *S. pneumoniae* isolates, serotype 33B | | China (Institute of Microbiology Chinese Academy of Sciences) |
| TH16810 | *S. pneumoniae* isolates, serotype 33C | | China (Institute of Microbiology Chinese Academy of Sciences) |
| TH16812 | *S. pneumoniae* isolates, serotype 34 | | China (Institute of Microbiology Chinese Academy of Sciences) |
| TH16815 | *S. pneumoniae* isolates, serotype 35A | | China (Institute of Microbiology Chinese Academy of Sciences) |
| TH16820 | *S. pneumoniae* isolates, serotype 35C | | China (Institute of Microbiology Chinese Academy of Sciences) |
| TH16821 | *S. pneumoniae* isolates, serotype 35F | | China (Institute of Microbiology Chinese Academy of Sciences) |
| TH16824 | *S. pneumoniae* isolates, serotype 37 | | China (Institute of Microbiology Chinese Academy of Sciences) |
| TH16827 | *S. pneumoniae* isolates, serotype 39 | | China (Institute of Microbiology Chinese Academy of Sciences) |
| TH16828 | *S. pneumoniae* isolates, serotype 41A | | China (Institute of Microbiology Chinese Academy of Sciences) |
| TH16829 | *S. pneumoniae* isolates, serotype 41F | | China (Institute of Microbiology Chinese Academy of Sciences) |
| TH16830 | *S. pneumoniae* isolates, serotype 48 | | China (Institute of Microbiology Chinese Academy of Sciences) |
| TH16920 | *S. pneumoniae* isolates, serotype 9A | | China (Peking Union Medical College Hospital) |
| TH16924 | *S. pneumoniae* isolates, serotype 24F | | China (Peking Union Medical College Hospital) |
| TH16926 | *S. pneumoniae* isolates, serotype 25A | | China (Peking Union Medical College Hospital) |
| TH16927 | *S. pneumoniae* isolates, serotype 25F | | China (Peking Union Medical College Hospital) |
| TH17104 | *S. pneumoniae* isolates, serotype 19B | | China (Institute of Microbiology Chinese Academy of Sciences) |

**Table S4. Bacterial strains and derivatives used in this study (Continued)**

| **Strain** | **Description** | **Reference or source** |
| --- | --- | --- |
| TH17251 | *S. pneumoniae* isolates, serotype 42 | China (Beijing Children's Hospita) |
| TH17797 | *S. pneumoniae* isolates, serotype 9L | China (Beijing Children's Hospita) |
| TH17798 | *S. pneumoniae* isolates, serotype 19C | China (Beijing Children's Hospita) |
| TH7338 | *S. pneumoniae* TH2740 derivative; TH2740∆*cps*::JC; Kan^R^ | This study |
| TH13133 | TH870 derivative; capsule switched strain; TH870∆*cps*::*cps*9V | (An et al., 2022) |
| TH14188 | TH870 derivative; capsule switched strain; TH870∆*cps*::*cps*19F | (An et al., 2022) |
| TH15937 | TH870 derivative; capsule switched strain; TH870∆*cps*::*cps*1 | (An et al., 2022) |
| TH15940 | TH870 derivative; capsule switched strain; TH870∆*cps*::*cps*4 | (An et al., 2022) |
| TH15942 | TH870 derivative; capsule switched strain; TH870∆*cps*::*cps*6B | (An et al., 2022) |
| TH15943 | TH870 derivative; capsule switched strain; TH870∆*cps*::*cps*8 | (An et al., 2022) |
| TH15944 | TH870 derivative; capsule switched strain; TH870∆*cps*::*cps*14 | (An et al., 2022) |
| TH17341 | TH2740 derivative; capsule switched strain; TH2740∆*cps*::*cps*12B | This study |
| TH17345 | TH2740 derivative; capsule switched strain; TH2740∆*cps*::*cps*12A | This study |
| TH17347 | TH2740 derivative; capsule switched strain; TH2740∆*cps*::*cps*11D | This study |
| TH17355 | TH2740 derivative; capsule switched strain; TH2740∆*cps*::*cps*10C | This study |
| TH17356 | TH2740 derivative; capsule switched strain; TH2740∆*cps*::*cps*10F | This study |
| TH17357 | TH2740 derivative; capsule switched strain; TH2740∆*cps*::*cps*24B | This study |
| TH17358 | TH2740 derivative; capsule switched strain; TH2740∆*cps*::*cps*36 | This study |
| TH17359 | TH2740 derivative; capsule switched strain; TH2740∆*cps*::*cps*40 | This study |
| TH17360 | TH2740 derivative; capsule switched strain; TH2740∆*cps*::*cps*46 | This study |
| TH17361 | TH2740 derivative; capsule switched strain; TH2740∆*cps*::*cps*47A | This study |
| TH17362 | TH2740 derivative; capsule switched strain; TH2740∆*cps*::*cps*47F | This study |
| TH17365 | TH2740 derivative; capsule switched strain; TH2740∆*cps*::*cps*32F | This study |
| TH17366 | TH2740 derivative; capsule switched strain; TH2740∆*cps*::*cps*33A | This study |
| TH17367 | TH2740 derivative; capsule switched strain; TH2740∆*cps*::*cps*33D | This study |

**Table S4. Bacterial strains and derivatives used in this study (Continued)**

| **Strain** | **Description** | | **Reference or source** |
| --- | --- | --- | --- |
| TH17228 | *Haemophilus influenzae* isolates, serotype a | Unknown | |
| TH17189 | *H. influenzae* isolates, serotype b | USA | |
| TH17229 | *H. influenzae* isolates, serotype c | USA | |
| TH17231 | *H. influenzae* isolates, serotype d | Netherlands | |
| NCTC10479 / TH17230 | *H. influenzae* isolates, serotype e | NCTC | |
| TH17232 | *H. influenzae* isolates, serotype f | China | |
| NTHi / TH17233 | *H. influenzae* isolates, non-typeable | Australia (Westmead Hospital, NSW) | |
| TH14157 | *Escherichia coli* isolates, serotype K1 | China (PLA 302 Hospital) | |
| TH14159 | *E. coli* isolates, serotype K4 | China (PLA 302 Hospital) | |
| TH14508 | *E. coli* isolates, serotype KG2-1 | China (Beijing Tsinghua Changgung Hospital) | |
| TH14510 | *E. coli* isolates, serotype KG2-3 | China (Beijing Tsinghua Changgung Hospital) | |
| DH5α | *E. coli* DH5α | | NEB |
| TH17167 | *E. coli* DH5α carrying plasmid pCMV-chikv-strepII with cDNA of murine *Fcna*. | | This study |
| TH17168 | *E. coli* DH5α carrying plasmid pCMV-chikv-strepII with cDNA of human *Fcn2*. | | This study |
| TH17169 | *E. coli* DH5α carrying plasmid pCMV-chikv-strepII with variant human *Fcn2*. Amino acids T75-Y97 of human FCN2 were replaced by R104-H126 of murine FCNA (hFcn*-*mF1). | | This study |
| TH17170 | *E. coli* DH5α carrying plasmid pCMV-chikv-strepII with variant human *Fcn2*. Amino acids V128-G146 of human FCN2 were replaced by I157-T175 of murine FCNA (hFcn*-*mF2). | | This study |
| TH17171 | *E. coli* DH5α carrying plasmid pCMV-chikv-strepII with variant human *Fcn2*. Amino acids N154-S164 of human FCN2 were replaced by Y183-Q193 of murine FCNA (hFcn*-*mF3). | | This study |
| TH17172 | *E. coli* DH5α carrying plasmid pCMV-chikv-strepII with variant human *Fcn2*. Amino acids V171-K186 of human FCN2 were replaced by Q200-Q215 of murine FCNA (hFcn*-*mF4). | | This study |
| TH17173 | *E. coli* DH5α carrying plasmid pCMV-chikv-strepII with variant human *Fcn2*. Amino acids A188-V202 of human FCN2 were replaced by S217-L231 of murine FCNA (hFcn*-*mF5). | | This study |
| TH17174 | *E. coli* DH5α carrying plasmid pCMV-chikv-strepII with variant human *Fcn2*. Amino acids S205-K221 of human FCN2 were replaced by T234-H250 of murine FCNA (hFcn*-*mF6). | | This study |

**Table S4. Bacterial strains and derivatives used in this study (Continued)**

| **Strain** | **Description** | **Reference or source** |
| --- | --- | --- |
| TH17175 | *E. coli* DH5α carrying plasmid pCMV-chikv-strepII with variant human *Fcn2*. Amino acids L227-V247 of human FCN2 were replaced by A256-Q276 of murine FCNA (hFcn*-*mF7). | This study |
| TH17176 | *E. coli* DH5α carrying plasmid pCMV-chikv-strepII with variant human *Fcn2*. Amino acids R256-N275 of human FCN2 were replaced by S285-H304 of murine FCNA (hFcn*-*mF8). | This study |
| TH17177 | *E. coli* DH5α carrying plasmid pCMV-chikv-strepII variant human *Fcn2*. The whole C-terminal fibrinogen-like domain (T75-A288 aa) of human FCN2 were replaced by R104-S317 of murine FCNA (hFcn*-*mFBG). | This study |
| TH17349 | *E. coli* DH5α carrying plasmid pcDNA3.4 with variant *Fcna*. The whole C-terminal fibrinogen-like domain (T100-S317 aa) of murine FCNA were replaced by Q71-A288 of human FCN2 (mFcn*-*hFBG). | This study |

Kan^R^: Kanamycin resistant; NEB: New England Biolabs (Beijing) LTD; NCTC: National Collection of Type Cultures
