## Supplementary material for "Plasma Ficolins Enable Liver Macrophages to Capture Blood-Borne Bacteria by Recognizing Capsular Polysaccharides": Table S5

**Table S5. Primers used in this study**

| **Primer ID** | **Sequence (5’-3’)** | **Description** |
| --- | --- | --- |
| Pr9840 | GAGATCTAGAGGATAATGCTGAAAACTCCTTGAAG | Forward primer to amplify Janus Cassette |
| Pr9199 | GAGACTCGAGCACAGAGTTTGTAGAAACGCAA | Reverse primer to amplify Janus Cassette |
| Pr19874 | GCAATGCCAGACAGTAACCTCTAT | Forward primer to amplify *cps*B gene for capsule type identification |
| Pr19875 | CCTGCCTGCAAGTCTTGATT | Reverse primer to amplify *cps*B gene for capsule type identification |
| Pr10489 | CAGTTTGAACATATCGGTCTTCAGT | Forward primer to amplify upstream homologous arm of *cps* locus from TH2740 |
| Pr10490 | GCTCTAGATCTGTCTTTTATACAGTCCTCCCTT | Reverse primer to amplify upstream homologous arm of *cps* locus from TH2740 |
| Pr10491 | CCGCTCGAGATGTTGTTTGAAAAATAATTTTC | Forward primer to amplify downstream homologous arm of *cps* locus from TH2740 |
| Pr10492 | GAACTGTCTGATCATCCAAAGCCT | Reverse primer to amplify downstream homologous arm of *cps* locus from TH2740 |
| Pr12134 | AGTGGATATCAATTACTATGTGCGAGGATAATGCTGAAAACTCCTTGAAG | Forward primer to amplify Janus Cssette (JC) |
| Pr12135 | GTTATTTCACATGTTTTGCGAGATCCAGAGTTTGTAGAAACGCAAAAAGG | Reverse primer to amplify Janus Cssette (JC) |
| Pr10704 | TGGTGTGACGTCCCCACATAATCAT | Forward primer to amplify *cps* locus |
| Pr10705 | TTATTGCGACCTCAAATGCTGGATT | Forward primer to amplify *cps* locus |
| Pr10710 | CTGGAGTCCATTTATAGGTGCTACCGTT | Reverse primer to amplify *cps* locus |
| Pr10711 | TGTTTTCAAAGGTACGAATTTCATCAAAGT | Reverse primer to amplify *cps* locus |
| Pr18850 | ACCCTGTGCCAGAGAGGACC | Forward primer to amplify FBG domain of *Fcna* from mouse liver cDNA |
| Pr20053 | AGATGCTCGGATTTTCATCTCGG | Reverse primer to amplify FBG domain of *Fcna* from mouse liver cDNA |
| Pr18904 | CTGACACGGGGCATCTTCCTGACTGGCTGGTACACCATCCATCTGCCCGACTGCCGGC | Forward primer containing R1 sequence of *Fcna* to linearize the plasmid pFCN-2 |
| Pr18905 | CAAGTCTTTGCAGCTCCGGGGTCCTCTCAGGCACGGCTGGGGCT | Reverse primer containing R1 sequence of *Fcna* to linearize the plasmid pFCN-2 |
| Pr18906 | ATAAAAGAGGCTTTGGCAACCTGGGCACGGAGTTCTGGCTGGGGAAT | Forward primer containing R2 sequence of *Fcna* to linearize the plasmid pFCN-2 |
| Pr18907 | AGGAGTCCCAGTCTCGGAAGAAATCGATAGAGCCATCCACCCTCCGCT | Reverse primer containing R2 sequence of *Fcna* to linearize the plasmid pFCN-2 |
| Pr18908 | GGAACCAAGAGCTCCGAGTTGACTTAGTGGACTTTGAGGACAACT | Forward primer containing R3 sequence of *Fcna* to linearize the plasmid pFCN-2 |
| Pr18909 | CATTGGCTGTGAGCAGGTGCAGGTAGTCATTCCCCAGCCAGAACT | Reverse primer containing R3 sequence of *Fcna* to linearize the plasmid pFCN-2 |
| Pr18910 | TATGCCAAGTACAGCTCATTCCAGGTAGCCGACGAGGCGGAGAAGT | Forward primer containing R4 sequence of *Fcna* to linearize the plasmid pFCN-2 |
| Pr18911 | GGAGCCTTTCCCTTGGAAATCTTGCAGGTCTACACGGAGCTCGCT | Reverse primer containing R4 sequence of *Fcna* to linearize the plasmid pFCN-2 |
| Pr18912 | TTGGGGCAGTTTCTGGAGGGCAGTGCGGGAGATTCCCTGAC | Forward primer containing R5 sequence of *Fcna* to linearize the plasmid pFCN-2 |
| **Table S5. Primers used in this study (Continued)** | | |
| **Primer ID** | **Sequence (5’-3’)** | **Description** |
| Pr18913 | GGTCAGCTTGTATTTCTCCTGTTCTTCAGACACCTTGAATGATCTGTACT | Reverse primer containing R5 sequence of *Fcna* to linearize the plasmid pFCN-2 |
| Pr19014 | CACAACAACATGTCATTTACAACCCATGACCAGGACAATGATC | Forward primer containing R6 sequence of *Fcna* to linearize the plasmid pFCN-2 |
| Pr19015 | CTTTGTCAGGGAGTCTCCTGCAGTGCCCTCCACGAAGG | Reverse primer containing R6 sequence of *Fcna* to linearize the plasmid pFCN-2 |
| Pr20324 | TGGAGCCTGGTGGTACCACAACTGCCACCAGTCCAACCTGAATGGTCGCTAC | Forward primer containing R7 sequence of *Fcna* to linearize the plasmid pFCN-2 |
| Pr20325 | ATTCATGCTGTTGGCATCATTGTCCTGGTCTTTGGTG | Reverse primer containing R7 sequence of *Fcna* to linearize the plasmid pFCN-2 |
| Pr20326 | CAACTGGGGAACTGGCCAAGGTCACCACTACTCCTACAAGGTGTCAGAG | Forward primer containing R8 sequence of *Fcna* to linearize the plasmid pFCN-2 |
| Pr20327 | ATGCCATCCGCATAACTCTCATGGCTCCCGCTGAGGTAGCGACCATTCAGG | Reverse primer containing R8 sequence of *Fcna* to linearize the plasmid pFCN-2 |
| Pr20330 | ATTCCCGCCGCCACCATGCAGTGGCCTACGCTGTG | Forward primer to amplify N-terminal sequence of *Fcna* from mouse liver cDNA |
| Pr19883 | GTCTCCCAGCTCCTTTTCACC | Reverse primer to amplify N-terminal sequence of *Fcna* from mouse liver cDNA |
| Pr20333 | ATTCCCGCCGCCACCATGGAGCTGGACAGAG | Forward primer to amplify N-terminal sequence of *Fcn2* from pFCN-2 |
| Pr20334 | TCTCTGGCACAGGGTGGGCTCCCCAGGTGCTCCG | Reverse primer to amplify N-terminal sequence of *Fcn2* from pFCN-2 |
| Pr19884 | AAGGAGCTGGGAGACCAGCCGTGCCTGACAGGCCCG | Forward primer to amplify FBG domain of *Fcn2* from pFCN-2 |
| Pr20331 | CCCTTAAGCTTATCATTTTTCGAACTGCGGGTGGC | Reverse primer to amplify FBG domain of *Fcn2* from pFCN-2 |
| Pr20048 | ATAAGAATGCGGCCGCGCCACCATGCAGTGGCCTACGCTGTGGGCCT | Forward primer to amplify murine *Fcna* from liver cDNA |
| Pr20049 | CGGGATCCTTATTTTTCGAACTGCGGGTGGCTCCACGATCCACCTCCAGATGCTCGGATTTTCATCTCGGCA | Reverse primer to amplify murine *Fcna* from liver cDNA |
| Pr20328 | TGATAAGCTTAAGGGTTCGATCCCT | Forward primer to linearize plasmid pcDNA3.4 |
| Pr20329 | GGTGGCGGCGGGAATTCAAGGG | Reverse primer to linearize plasmid pcDNA3.4 |
| Pr20332 | AAAATCCGAGCATCTGGAGGTGGATCGTGGAGCCAC | Forward primer to linearize plasmid pcDNA3.4 |
| Pr20323 | GGTGGCGGCGGGAATTCAAG | Reverse primer to linearize plasmid pcDNA3.4 |
