## Supplementary material for "Plasma Ficolins Enable Liver Macrophages to Capture Blood-Borne Bacteria by Recognizing Capsular Polysaccharides": Table S6

**Table S6.** **Information for constructions of strains used in this study**

| **Strain ID** | **Genotype** | **Donor DNA** | **Template**  **DNA** | **Digestion and ligation** | **Recipient or parental strain** |
| --- | --- | --- | --- | --- | --- |
| TH7338 | TH2740∆*cps*::JC | Pr10489/10490^a^  Pr10491/10492^b^  Pr9840/9199^c^ | TH2740/JC | XbaI/XhoI | TH2740 |
| TH17345 | TH2740∆*cps*::*cps*12A | Pr10705/10711^c^ | Genomic DNA with *cps*12A locus^d^ | ／ | TH7338 |
| TH17347 | TH2740∆*cps*::*cps*11D | Pr10705/10711^c^ | Genomic DNA with *cps*11D locus^d^ | ／ | TH7338 |
| TH17355 | TH2740∆*cps*::*cps*10C | Pr10705/10711^c^ | Genomic DNA with *cps*10C locus^d^ | ／ | TH7338 |
| TH17356 | TH2740∆*cps*::*cps*10F | Pr10705/10711^c^ | Genomic DNA with *cps*10F locus^d^ | ／ | TH7338 |
| TH17357 | TH2740∆*cps*::*cps*24B | Pr10705/10711^c^ | Genomic DNA with *cps*24B locus^d^ | ／ | TH7338 |
| TH17358 | TH2740∆*cps*::*cps*36 | Pr10705/10711^c^ | Genomic DNA with *cps*36 locus^d^ | ／ | TH7338 |
| TH17359 | TH2740∆*cps*::*cps*40 | Pr10705/10711^c^ | Genomic DNA with *cps*40 locus^d^ | ／ | TH7338 |
| TH17360 | TH2740∆*cps*::*cps*46 | Pr10705/10711^c^ | Genomic DNA with *cps*46 locus^d^ | ／ | TH7338 |
| TH17361 | TH2740∆*cps*::*cps*47A | Pr10705/10711^c^ | Genomic DNA with *cps*47A locus^d^ | ／ | TH7338 |
| TH17362 | TH2740∆*cps*::*cps*47F | Pr10705/10711^c^ | Genomic DNA with *cps*47F locus^d^ | ／ | TH7338 |
| TH17365 | TH2740∆*cps*::*cps*32F | Pr10704/10710^c^ | Genomic DNA with *cps*32F locus^d^ | ／ | TH7338 |
| TH17366 | TH2740∆*cps*::*cps*33A | Pr10705/10711^c^ | Genomic DNA with *cps*33A locus^d^ | ／ | TH7338 |
| TH17367 | TH2740∆*cps*::*cps*33D | Pr10705/10711^c^ | Genomic DNA with *cps*33D locus^d^ | ／ | TH7338 |

^a^ Primers used to amplify upstream homologous arm from recipient strain

^b^ Primers used to amplify downstream homologous arm from recipient strain

^c^ Primers used to amplify insertion fragment

^d^ Genomic DNA carrying different types of capsule synthesis loci
